# Surviving phage attack primes bacterial immunity to defeat counterdefenses

**DOI:** 10.1101/2025.11.13.688357

**Authors:** Reid T. Oshiro, Alyssa A. Pratt, Drew T. Dunham, Lindsey A. Doyle, Clare Gill, Ani Chouldjian, Kerilyn M. Higashi, Julieta Peralta Acosta, Sydney F. McGuire, Sam Nouwens, Brett K. Kaiser, Kimberley D. Seed

## Abstract

Transcriptional repressors with ligand-responsive WYL domains control diverse bacterial defense systems, yet the natural cues driving derepression and the physiological rationale for dynamic regulation in native host-phage contexts remain unknown. Here, we show that restricted phage infection, where defense clears a primary attack, serves as a natural cue for WYL-mediated derepression in *Vibrio cholerae*. Although not required to block the initial infection, this response increases defense protein abundance, shifting host-phage stoichiometry and priming surviving bacteria to overwhelm phage-encoded counterdefenses in subsequent attacks. We also find that restricted infection co-induces an unlinked anti-plasmid defense via its own WYL repressor, showing that parallel WYL sensors coordinate a broader immune response without regulatory crosstalk. In contrast, productive phage infection triggers horizontal transfer of the defense-encoding mobile element, ensuring its persistence in the population. Together, our work reveals that infection fate dictates divergent outcomes for the expression and dissemination of bacterial immunity.

## Introduction

Bacteria have evolved a diverse and rapidly expanding arsenal of defense systems, ranging from single-protein effectors to multi-protein complexes, that sense and restrict bacterial viruses, or phages^1–5^. With over 200 distinct systems identified^3,4,6^, these defenses employ a range of mechanisms to hinder phage replication, from the direct nucleolytic cleavage of phage DNA by restriction-modification (RM)^7,8^ and CRISPR-Cas^9–12^ systems, to the induction of host cell death or dormancy by systems like CBASS^13^ and toxin-antitoxin^14^ (TA) systems to halt viral spread. While our mechanistic understanding of how these defense systems function has grown dramatically, a critical, yet poorly understood aspect of their biology is how their expression is regulated in their native hosts, particularly in response to phage infection.

Many phage defense systems are controlled by genetically-linked repressors that maintain low basal expression^15–19^. This repression is thought to mitigate toxicity or prevent cell death associated with spurious immune activity in the absence of infection^15,18^. Although it is hypothesized that this repression is alleviated upon phage challenge, whether this induction occurs and what specific protective advantage it confers remains ambiguous. A particularly widespread family of defense-associated repressors is characterized by the presence of a WYL domain (WYL proteins), and is associated with several prominent defense systems, including CRISPR-Cas, TAs, BREX, and CBASS^16,20,21^. These ligand-sensing transcriptional repressors have been shown repress expression of their linked systems by binding to inverted repeats in the linked defenses system’s promoter region^16–18^. Biochemical models suggest that binding to single-stranded DNA (ssDNA) can relieve this repression and upregulate defense^22^. However, prior research has focused on artificial induction or non-native systems, so the natural cues that lead to derepression—and the biological benefits of this regulation in native contexts— remain unknown.

The pairing of WYL proteins with phage defense systems is found within the SXT/R391 family of integrative and conjugative elements (ICEs)^21^. The carriage of defense systems on mobile genetic elements (MGEs), such as ICEs, is a common theme in bacterial immunity, enabling the rapid horizontal transfer of entire defensive arsenals between bacteria^23^. SXT ICEs encode both phage defense systems and antibiotic resistance genes^21,24,25^, a combination of significant clinical relevance. In clinical isolates of *Vibrio cholerae,* the causative agent of the diarrheal disease cholera, these ICEs influence the sensitivity to multiple phages, including ICP1, the predominant lytic phage found in cholera patient stools^26^. The most prevalent SXT ICE in *V. cholerae* is *Vch*Ind5, which encodes a Type I <u>B</u>acteriophage <u>Ex</u>clusion (BREX) system. Following our initial discovery of BREX’s regulation by a proximal WYL protein in *Vch*Ind5^21^, homologous repressors (now broadly termed BrxR) have been identified for numerous other BREX systems^16,17,21^. The *Vch*Ind5 ICE and its inhibition of the phage ICP1 provide an ideal model system to investigate the biological purpose of WYL-mediated regulation. Crucially, this system also includes a phage-encoded counterdefense protein, OrbA, that inhibits the activity of the *Vch*Ind5-encoded BREX system^21,27^. Together, this complete set of interactions between a host bacterium, its endogenous defense system and repressor, and a phage with a cognate counterdefense provides a unique opportunity to elucidate the biological importance of phage defense regulation and to assess how different phage infection outcomes impact immune system expression.

Here, we leverage the native *V. cholerae* BrxR-BREX system to determine how two distinct phage infection outcomes impact defense expression. We find that a restricted infection, in which the host cell survives, induces BREX expression through BrxR-mediated sensing of ssDNA, whereas generic DNA damage does not recapitulate the same response, suggesting a phage-specific signal. In contrast, a productive infection, resulting in host cell lysis and release of phage progeny, triggers a divergent response, inducing the horizontal transfer of the mobile element carrying the defense system. Rather than serving to prevent severe fitness costs, dynamic BREX derepression functions as an immune priming mechanism that proactively shifts defense-counterdefense stoichiometry, protecting surviving bacteria against subsequent attacks by evolved phages encoding counterdefenses like OrbA. Furthermore, we find that restricted infection simultaneously co-induces an unlinked anti-plasmid defense system (DdmDE) through its own WYL repressor, demonstrating that independent WYL proteins can coordinate the upregulation of distinct defenses in response to a neutralized phage threat. Bioinformatic analyses reveal that co-occurring defense-associated WYL regulators are widespread across bacterial taxa, suggesting that independent WYL transcription factors operating in parallel to coordinate multi-layered immune responses represent a common feature of bacterial immunity. Collectively, our findings establish a broadly applicable, physiologically relevant paradigm for WYL-mediated defense regulation. Further, these findings reveal a striking, coordinated intracellular regulatory response in bacteria conceptually analogous to the intercellular response in vertebrate innate immunity, wherein viral infection triggers a cascade of immune signaling that amplifies the host protection.

## Results

### BREX expression increases during a restricted phage infection, while SXT transfer gene expression increases during productive phage infection

To resolve how different phage infection outcomes shape the transcriptional landscape of a mobile genetic element and its encoded defense system, we performed RNA sequencing. We modeled a productive infection, where the phage bypasses host defense, by infecting BREX(+) cells with an ICP1 phage encoding the BREX-counterdefense OrbA (ICP1). To model a restricted infection, where host defense is active and effective, we used an isogenic phage with OrbA deleted (ICP1 Δ*orbA*). Since BREX inhibits ICP1 Δ*orbA* from initiating replication^27^, samples were collected immediately following the addition of phage (0 minutes), before the start of replication (4 minutes post-infection), and after the start of theta replication^28^ (10 minutes post-infection). Differential expression analysis using DESeq2^29^ comparing the restricted and productive infection transcriptomes revealed the specific upregulation of BREX genes (*brxRABCXZL*) at the 4- and 10-minute time points during the restricted infection (Fig. 1A). Serendipitously, we also observed upregulation of the genetically unlinked anti-plasmid defense system DdmDE^30^ during restricted infection, which is hypothesized to be regulated by an upstream WYL protein^31^. In contrast, SXT transfer genes, which are required for the element’s horizontal spread via conjugation^24,32,33^, were downregulated in this same comparison by 10 minutes post-infection.

**Figure 1.**
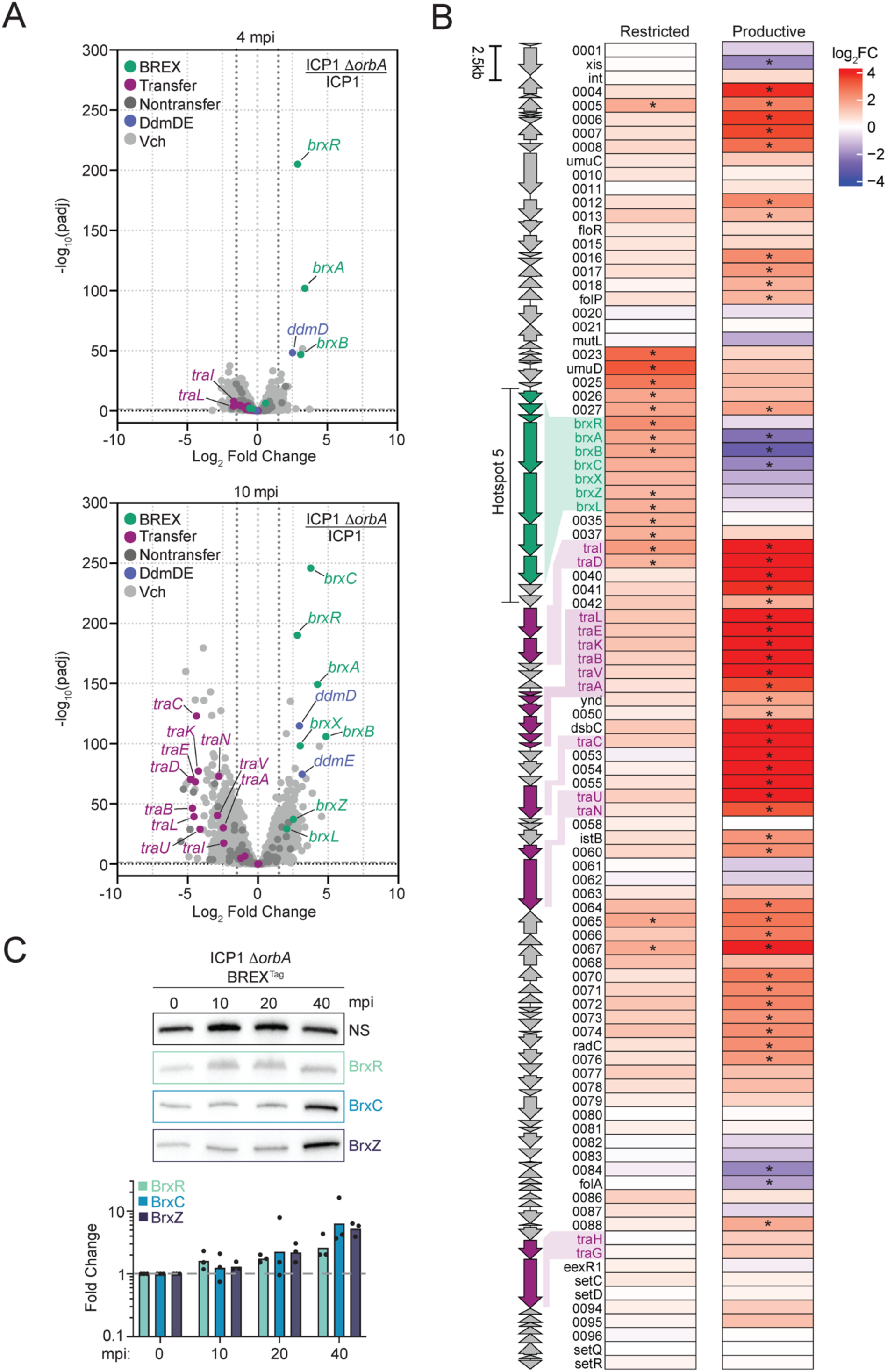
Restricted phage infection induces BREX expression. **A)** Volcano plots showing changes in gene expression, plotted as the Log_2_ fold change of a restricted (ICP1 Δ*orbA*) infection relative to the productive (ICP1) phage infection at 4 minutes (top) and 10-minutes (bottom) post-infection (mpi). Differential expression was determined using DESeq2^29^. SXT transfer (*tra*) genes were assigned based on the following references^24,32,33^. Genes encoded by SXT that are not involved in SXT transfer or BREX are categorized as nontransfer. Vch denotes genes encoded in the *V. cholerae* chromosome outside of the SXT ICE. −Log_10_ (p_adj_) ≥ 1.25 and Log_2_ fold change ≥ ± 1.5 were considered significant (grey dashed and dotted lines, respectively). See Supplementary Tables 1 and 2 for exact values. **B)** Heat map displaying the Log_2_ fold change across the SXT ICE *Vch*Ind5 element during a restricted (left) or productive (right) infection, comparing expression at 10 mpi to their respective 0-minute time point. Next to the heat maps is a schematic of the genomic architecture of the SXT ICE element. * −Log_2_ fold change ≤ −1.5 or ≥ 1.5 with a p_adj_ value < 0.05. SXT transfer genes were assigned as the same in panel A. See Supplementary Tables 3 and 4 for exact values. **C)** Western blot analysis of BREX protein abundance at different time points during a restricted phage infection using the tagged BREX strain (BREX^Tag^). The primary antibody against BrxR detects both BrxR and a non-specific (NS) band, which was used as a loading control. Top: A representative biological replicate; for additional replicates, see Supplementary Figure 2. Below: Quantification of the indicated proteins during ICP1 Δ*orbA* infection at different time points. Fold change was calculated by normalizing each protein’s intensity to the NS loading control. Dots represent the individual biological replicates, and bars represent the mean. Statistical analysis was performed using a One-way ANOVA with Dunnett’s multiple-comparison test comparing each protein quantification to its respective zero-minute time point, and the results were not significant (*P* > 0.05).

To assess whether differences in SXT ICE gene expression between restricted and productive infection reflect active induction or repression, we compared the 10-minute time point for each infection to its respective baseline (t = 0 min; Fig. 1B). During restricted phage infection, overall transcriptional changes across the element were limited, with only 17 of 83 SXT ICE genes, including 5 of 7 BREX genes, significantly upregulated, and none downregulated (Fig. 1B). In contrast to the relative downregulation seen when comparing infection outcomes directly, SXT transfer genes were not actively downregulated during restricted infection; rather, core transfer machinery remained largely unchanged, with only 2 of 14 transfer genes showing significant upregulation—both located directly downstream of the BREX locus (Supplementary Table 3). Conversely, by 10 minutes post-infection, a productive infection elicited a strikingly divergent response across the element, with 53 of 83 genes differentially regulated (Fig. 1B). This response was driven by the robust upregulation of 47 genes— including 12 of 14 transfer genes—and the downregulation of 6 genes, including 3 BREX genes. This rapid induction of transfer machinery within minutes of infection provides a transcriptional basis for our previous observation of increased SXT conjugation during productive phage infection, a process previously assayed only after several hours^21^. Together, these baseline comparisons demonstrate that the SXT ICE *Vch*Ind5 mounts divergent transcriptional responses to phage attack: restricted infection induces the element’s defense system, while productive infection triggers its horizontal transfer.

To determine whether the transcriptional upregulation of BREX translates into increased protein levels, we monitored the abundance of two components of the BREX effector complex^34^ alongside their WYL-domain regulator, BrxR. We engineered a strain (BREX^Tag^) in which complex components BrxC and BrxZ were tagged at their native loci with 3x FLAG and HA, respectively, while BrxR was detected using a custom antibody (Supplementary Fig. 1). BrxC was selected for its essential role in BREX function^27^ and its interaction with OrbA^27^, whereas BrxZ served as a measure for the downstream portion of the BREX locus. After confirming the BREX^Tag^ strain phenocopied the wild-type’s defense activity (Extended Data Fig. 1), we challenged it with ICP1 Δ*orbA*. The observed transcriptional induction was followed by a corresponding increase in BREX protein abundance, which became apparent 40 minutes post-infection (Fig. 1C). Densitometry analysis revealed a ∼2-fold increase in BrxR, while BrxC and BrxZ increased more dramatically, by ∼5- and 4-fold, respectively (Fig. 1C). This modest increase in BrxR compared to the effector components may reflect autoregulatory or post-transcriptional mechanisms to minimize enhanced repression following phage clearance. The delay in protein accumulation relative to transcription likely reflects the time required for translation during active phage restriction. Together, these results confirm that transcriptional induction during restricted phage infection is accompanied by increased defense protein abundance.

### Structural basis for BrxR derepression by single-stranded DNA during restricted phage infection

To confirm that BrxR mediates the increase in BREX expression during restricted phage infection (Fig. 1), we measured BREX protein levels during ICP1 Δ*orbA* infection in a *brxR*-deficient background (BREX^tag^ Δ*brxR*). Unlike the robust protein induction observed in wild-type cells (Fig. 1C), BrxC and BrxZ levels remained unchanged in the absence of BrxR (Fig. 2A). These results show that BrxR is essential for dynamic BREX system expression during restricted phage infection.

**Figure 2.**
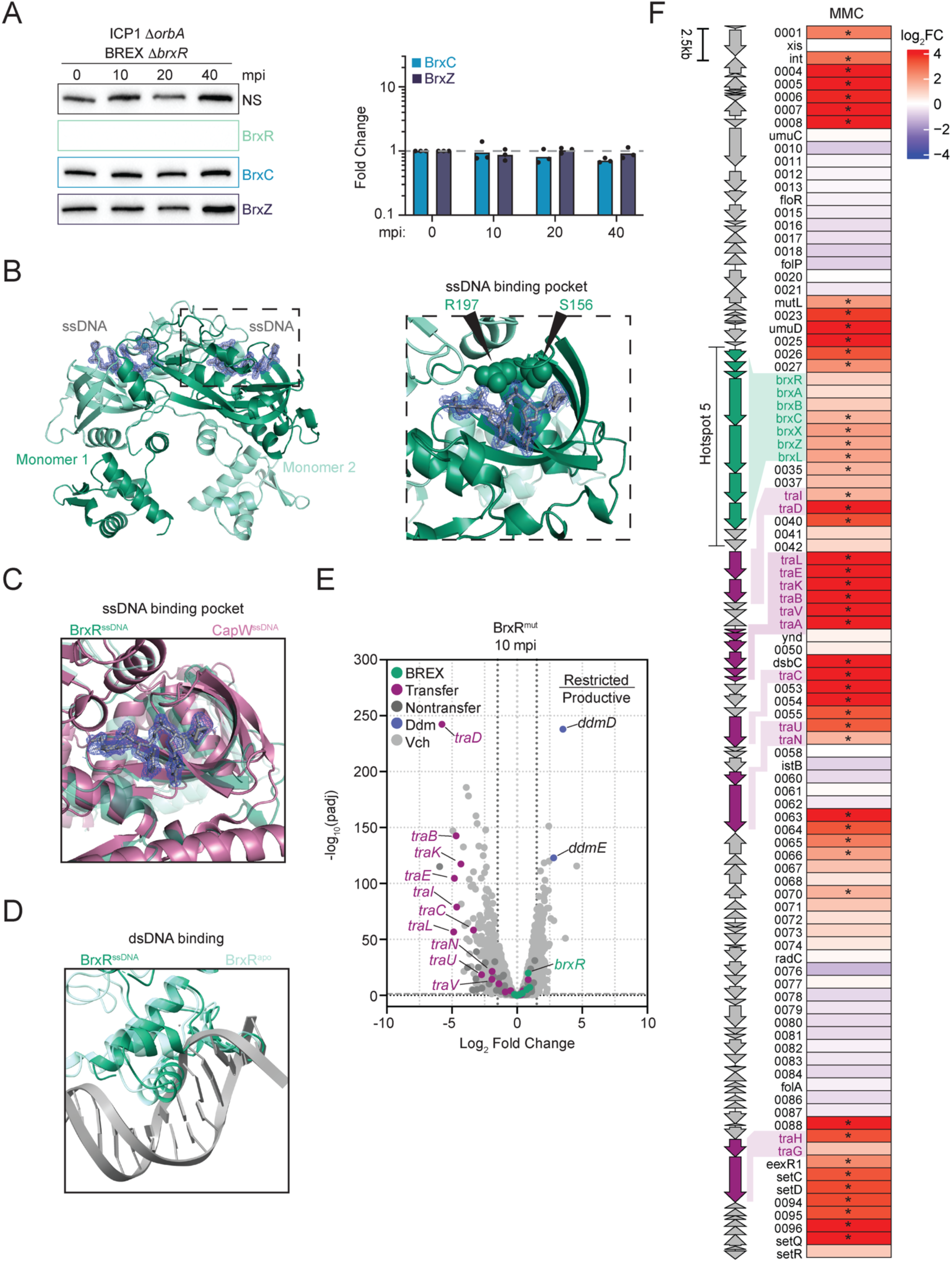
Interaction with single-stranded DNA regulates BrxR repressor activity. **A)** Western blot analysis of BREX protein abundance at different time points during restricted phage infection using the BrxR-deletion-tagged BREX strain (BREX^Tag^ Δ*brxR*). Left: A representative biological replicate; for additional replicates, see Supplementary Figure 2. Right: Quantification of the indicated proteins during ICP1 Δ*orbA* infection at different time points. Dots represent the individual biological replicates, and bars represent the mean. Statistical analysis was performed using a One-way ANOVA with Dunnett’s multiple-comparison test comparing each protein quantification to its respective zero-minute time point, and the results were not significant (*P* > 0.05). **B)** 3D crystal structure of BrxR encoded by the SXT ICE *Vch*Ind5 in the presence of single-stranded (ss) DNA. The electron density of ssDNA is shown in the ssDNA binding pocket of BrxR. The inlet is a zoom in of the ssDNA-binding pocket, displaying two residues that contact the ssDNA. Crystallographic statistics are included in Supplementary Table 5. **C)** A zoom-in of the ssDNA-binding pocket of BrxR with the electron density of the ssDNA compared to the ssDNA-bound CapW structure (PDB: 9C5G); the CapW ssDNA is not present in the structure. **D)** A zoom-in on the winged helix-turn-helix domain of an overlay of the apo and ssDNA-bound structure of BrxR overlaid with the *Acinetobacter* BrxR structure bound to double-stranded (ds) DNA (PDB: 7T8K) to emphasize the clash that occurs when BrxR is bound to ssDNA. The Acinetobacter BrxR is not displayed. **E)** Volcano plots showing changes in gene expression in the BrxR^mut^ background, plotted as the Log_2_ fold change of the restricted (ICP1 Δ*orbA*) relative to the productive (ICP1) phage infection, at 10-minute post-infection (mpi). Differential expression was determined using DESeq2^29^. SXT transfer (*tra*) genes were assigned based on the following references^24,32,33^. Genes encoded by SXT that are not involved in SXT transfer or BREX are categorized as nontransfer. Vch denotes genes encoded in the *V. cholerae* chromosome outside of the SXT ICE. −Log_10_ (p_adj_) ≥ 1.25 and Log_2_ fold change ≥ ± 1.5 were considered significant (grey dashed and dotted lines, respectively). See Supplementary Table 6 for the exact values. **F)** Heat map displaying the Log_2_ fold change across the SXT ICE *Vch*Ind5 element after 60 minutes of treatment with mitomycin C compared to its untreated 0-minute time point. Next to the heat maps is a schematic of the genomic architecture of the SXT ICE element. * −Log_2_ fold change ≤ −1.5 or ≥ 1.5 with a p_adj_ value < 0.05. SXT transfer genes were assigned as the same as in Fig. 1. See Supplementary Table 7 for exact values.

The model for WYL-domain proteins as ligand-sensing repressors stems largely from studies of CapW, which regulates a CBASS defense system^35^. *In vitro*, single-stranded DNA (ssDNA) binding to CapW’s WYL domain induces a conformational change that releases the repressor from its promoter, activating transcription^22^. To test if Vch*Ind5* BrxR uses a similar mechanism, we solved crystal structures of BrxR with and without ssDNA (Fig. 2B and Extended Data Fig. 2A). Structural alignment of ssDNA-bound BrxR and CapW revealed striking conservation within their ssDNA binding pockets—contacting analogous residues despite sharing only 26% overall sequence identity (Fig. 2C and Extended Data Fig. 2B). As in CapW, ssDNA binding to BrxR causes an allosteric conformational change that separates its winged helix-turn-helix domains, creating steric clashes with the DNA backbone that force repressor dissociation (Fig. 2D and Extended Data Fig. 2C).

To test if BREX derepression during restricted phage infection requires BrxR’s ssDNA-binding interface, we engineered an ssDNA-binding mutant (BrxR^mut^) targeting key contact residues in the ssDNA-binding pocket (S156 and R197; Fig. 2B right). To validate promoter binding, we identified BrxR’s target inverted repeat in the BREX promoter region using EMBOSS palindrome^36^. Recombinant wild-type BrxR bound this sequence with

∼250 nM affinity, while binding was lost when the sequence was scrambled (Extended Data Fig. 3A-B). BrxR^mut^ retained double-stranded DNA (dsDNA) binding, though with a modest ∼2-fold reduction compared to wild-type (Extended Data Fig. 3C). However, while ssDNA competition efficiently displaced wild-type BrxR from its dsDNA target (Extended Data Fig. 3D), BrxR^mut^ remained resistant to ssDNA and stayed bound to the promoter dsDNA, confirming it is blind to ssDNA-mediated derepression (Extended Data Fig. 3E).

We next asked if ssDNA sensing by BrxR is required for BREX induction during restricted phage infection. We modeled both productive and restricted infections in the BrxR^mut^ background and compared their transcriptional profiles at 10 minutes post-infection. In BrxR^mut^, BREX induction during restricted infection was completely abolished, while *ddmDE* induction was unaffected (Fig. 2E), consistent with *ddmDE* being controlled by its own upstream WYL-domain regulator. These results demonstrate that BrxR’s ssDNA binding is necessary for BREX derepression during restricted phage infection.

To put our findings in context with other WYL-domain regulators such as CapW, which responds to DNA-damaging agents like mitomycin C (MMC), we evaluated the transcriptomic response of *V. cholerae* after 60 minutes of MMC treatment (Fig. 2F). Interestingly, MMC induced a distinct SXT ICE transcriptional profile that was different from both restricted and productive phage infection. MMC treatment drove strong induction of SXT ICE transfer genes, consistent with classic SOS-dependent mobilization^37,38^. However, BREX upregulation was mild and limited to downstream genes of the locus, with *brxR* expression remaining unchanged. These results indicate that while ssDNA binding is necessary for BrxR derepression, global ssDNA generated during SOS-inducing DNA damage is not sufficient for robust BREX activation. Instead, restricted phage infection appears to represent a unique physiological state that selectively maximizes defense operon induction without triggering full-scale element mobilization.

### Dynamic expression of BREX promotes increased protection against subsequent infections by phages encoding inhibitors

Having established that BrxR dynamically derepresses BREX during restricted phage infection, we next sought to determine the physiological role of this upregulation. To test whether BrxR-mediated BREX upregulation is necessary to clear the primary, defense-sensitive phage, we challenged the unresponsive BrxR^mut^ strain (Extended Data Fig. 4A) with ICP1 Δ*orbA*. Preventing BREX induction did not affect phage restriction efficiency compared to wild-type cells, indicating that baseline BREX levels are sufficient to clear defense-sensitive phage. This finding aligns with infection kinetics: because ICP1 completes its lytic cycle in ∼20 minutes, whereas BREX protein accumulates only by ∼40 minutes post-restriction (Fig. 1C), defense induction occurs too late to contribute to clearance of the primary infection, suggesting this induction functions downstream.

Viral counterdefenses often function by binding and inactivating defense complexes, making the stoichiometry between defense proteins and inhibitors critical for infection outcome^39–41^. In clinical settings, *V. cholerae* encounters dynamic populations of ICP1, with both OrbA-expressing and non-expressing variants^21^. We reasoned that BREX upregulation during a primary restricted infection might act as an immune priming mechanism, increasing defense protein levels to protect surviving cells from subsequent attacks by phages with counterdefenses.

To test this model, we first asked whether elevated BREX levels can overcome OrbA-mediated counterdefense. Cells with constitutively high BREX expression (Δ*brxR*; Fig. 3A) were challenged with wild-type OrbA-encoding ICP1 (OrbA^+^). High BREX levels reduced ICP1 plaquing efficiency by nearly 100-fold (Fig. 3B) and eliminated phage production in single-round infection assays (Fig. 3C). Deleting two downstream BREX locus genes, encoding a protein of unknown function and a predicted abortive infection protein, did not change this effect, confirming that enhanced protection was due to BREX (Extended Data Fig. 4B and 4C). Inducing BREX expression to ∼10-fold above wild-type levels with an inducible promoter further reduced ICP1 plaquing efficiency by ∼15,000-fold (Fig. 3D). Together, these results show that the stoichiometric ratio of BREX to OrbA determines infection outcome and that elevated BREX levels can overwhelm counterdefense-mediated evasion.

**Figure 3.**
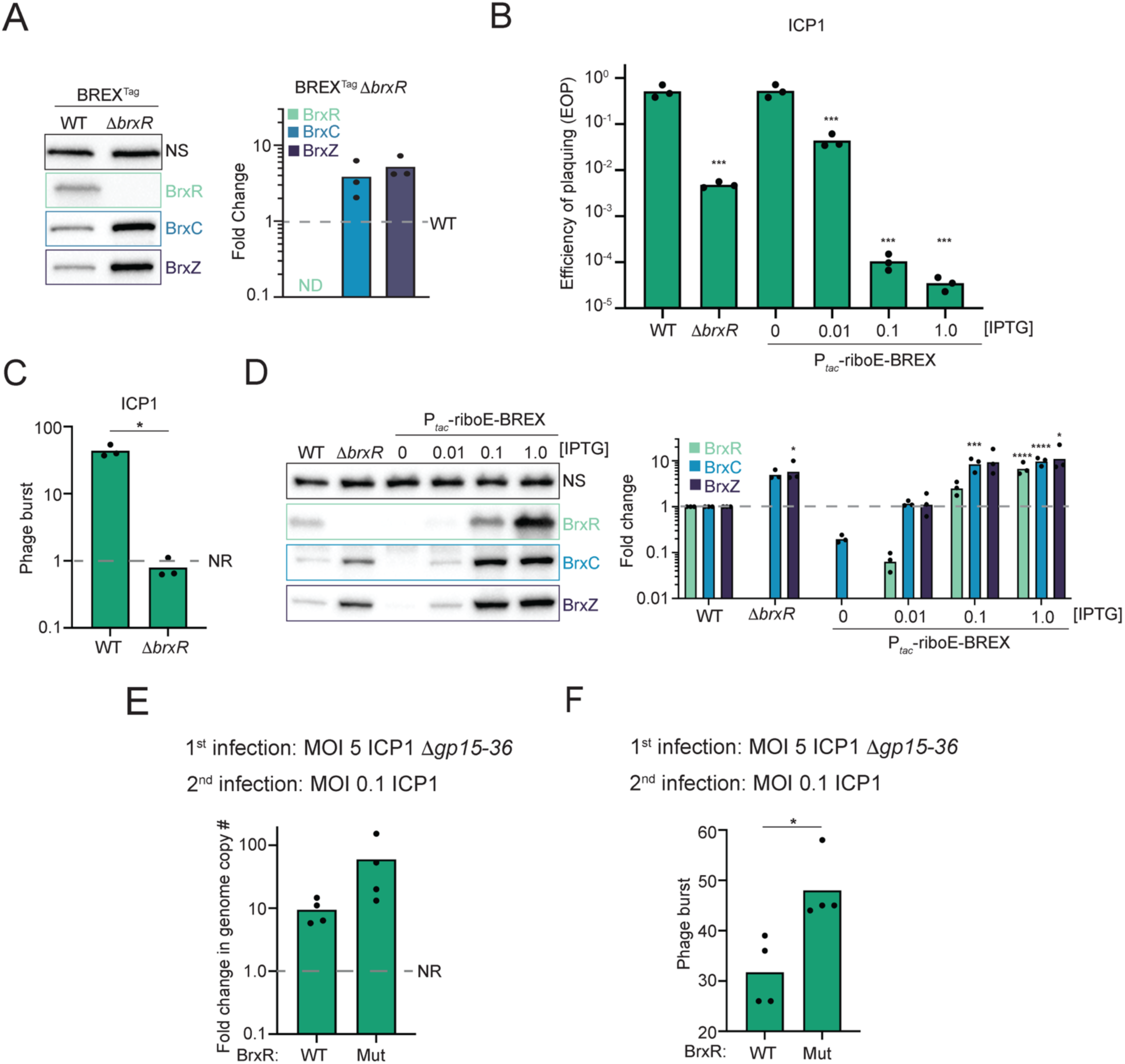
Restricted infection primes cells to stoichiometrically outcompete phage-encoded inhibitor to reduce phage progeny production. **A)** Western blot and densitometry analysis of BrxC and BrxZ in the absence of BrxR. For additional replicates and uncropped gels, see Supplementary Figure 5A. Fold change was calculated by normalizing each protein’s intensity to the NS loading control. Dots represent the individual biological replicates, and bars represent the mean. A Student’s t-test was performed for protein quantification, and the results were not significant (*P* > 0.05). **B)** Efficiency of plaquing of ICP1 in the presence and absence of BrxR and the inducible BREX background (*P_tac_*-*riboE*-BREX^tag^) using different inducer concentrations. DL indicates the detection limit. Statistical analysis was performed using a One-way ANOVA with Dunnett’s multiple-comparison test comparing each condition to wild type (*** = *P* < .001). **C)** Burst assay of ICP1 in the presence and absence of BrxR. The dashed line (NR = 1) indicates no net phage replication (output phage equals = input phage). Statistical analysis was performed using a Student’s t-test (* = *P <* 0.05). **D)** Western blot and densitometry analysis of BREX protein abundance at different inducer concentrations using the *P_tac_*-*riboE*-BREX^tag^ strain. For additional replicates, see Supplementary Fig. 5B. Fold change was calculated the same as in panel A. Dots represent the individual biological replicates, and bars represent the means. Statistical analysis was performed using a One-way ANOVA with Dunnett’s multiple-comparison test, comparing each protein to its respective wild-type condition (* = *P* < .05, *** = *P* < .001, **** = *P* < .0001). **E)** Quantification of ICP1 genome replication during a subsequent infection. Cells were first infected with ICP1 Δ*gp15-36*, and later infected with ICP1. The dashed line at one indicates there was no genome replication (NR). Each dot represents an individual biological replicate, and each bar represents the mean of the displayed biological replicates. Statistical analysis was performed using a Student’s t-test and was found not to be significantly different (*P* > 0.05). **F)** Burst assay of ICP1 during a subsequent infection. Cells were first infected with ICP1 Δ*gp15-36*, and later infected with ICP1. Burst size was enumerated on a BREX-positive strain. The dashed line (NR = 1) indicates no net phage replication (output phage equals = input phage). Statistical analysis was performed using a Student’s t-test (* = *P <* 0.05). Each dot represents an individual biological replicate.

To determine whether BREX upregulation during a primary restricted infection protects against secondary attack by counterdefense-encoding phages, we established a sequential infection assay. Cells were first primed by a high multiplicity of infection (MOI = 5) using defense-sensitive phage ICP1 Δ*orbA* to induce widespread BREX derepression across the population. Forty minutes post-restriction, a time point when BREX protein levels are elevated (Fig. 1C), primed cells were challenged with wild-type, OrbA-encoding ICP1, and secondary phage genome replication was measured by qPCR. Unexpectedly, wild-type ICP1 did not replicate in the wild-type background (Extended Data Fig. 4D), contrasting with our previous findings demonstrating phage replication during primary infection in the presence of BREX^27^. Replication also did not recover in the unresponsive BrxR^mut^ strain, suggesting this suppression occurred independently of BREX (Extended Data Fig. 4D).

Because lytic phages can encode early-expressed superinfection exclusion (SIE)^42^ factors that can block secondary infection by related phages^43,44^, we hypothesized that primary ICP1 infection could render cells refractory to subsequent infection, even when viral replication is blocked by host defense. To control for confounding SIE activity, we generated a phage deletion mutant lacking a large immediate-early genomic region (Δ*gp15-36*; see methods) to eliminate putative SIE factors. Using ICP1 Δ*gp15-36* to prime cells by restricted infection, we observed genome replication of secondary OrbA^+^ ICP1 in wild-type cells (Fig. 3E and Extended Data Fig. 4D), indicating the secondary phage bypassed SIE. Importantly, the derepression-deficient BrxR^mut^ background allowed modestly higher phage DNA replication, suggesting that cells unable to increase BREX expression during phage restriction are more permissive to secondary phage replication.

To evaluate whether this trend in reduced DNA replication impacts the complete viral lifecycle, we measured progeny virion production using burst assays after secondary challenge. Consistent with the replication data, phage production was significantly higher in the BrxR^mut^ background than in wild-type BrxR cells (Fig. 3F), indicating that failure to upregulate BREX compromises host defense against counterdefense-encoding phages. This assay likely underestimates the full extent of priming-mediated protection, as BrxR^mut^ has slightly higher basal BREX expression due to reduced dsDNA-binding affinity, resulting in a ∼2-fold decrease in OrbA+ ICP1 plaquing efficiency (Extended Data Fig. 3 and 4A). Collectively, these findings support a model in which primary restricted infection relieves BrxR repression, dynamically increasing BREX protein levels to protect surviving bacteria from subsequent attack by counterdefense-encoding phages.

### Constitutively high BREX expression alters the host transcriptome without a fitness cost or disruption to SXT ICE transfer efficiency

Having established that elevated BREX levels protect host cells against counterdefense-encoding phage, we next evaluated the physiological rationale for keeping the system repressed under baseline conditions. A prevailing hypothesis in the field is that defense repressors exist to minimize metabolic burden or prevent off-target toxicity prior to infection^45^. We therefore examined whether unregulated BREX expression imposes host fitness costs, alters global transcription or disrupts horizontal transfer of the defense system. To determine whether BrxR regulates host genes beyond the BREX system, we performed RNA sequencing on uninfected cells and compared the transcriptome of the Δ*brxR* mutant with that of the wild-type strain. As expected, derepression in the Δ*brxR* mutant resulted in significant upregulation of the BREX operon (Fig. 4A). However, we also observed that 224 host genes (5.56% of the genome) were differentially expressed, reflecting broad transcriptomic perturbations characterized by the downregulation of central metabolism and upregulation of cofactor synthesis pathways (Extended Data Fig. 5A and B). To distinguish whether BrxR functions as a global regulator or whether these effects were indirect, we next overexpressed BrxR from a neutral chromosomal locus. In this context, only the BREX system was significantly downregulated (Fig. 4B). Notably, *ddmDE*, which was also upregulated with BREX during restricted infection (Fig. 1A), was not regulated by BrxR in either the deletion or overexpression datasets (Fig. 4A and B), consistent with *ddmDE* being controlled by its own upstream WYL-domain regulator.

**Figure 4.**
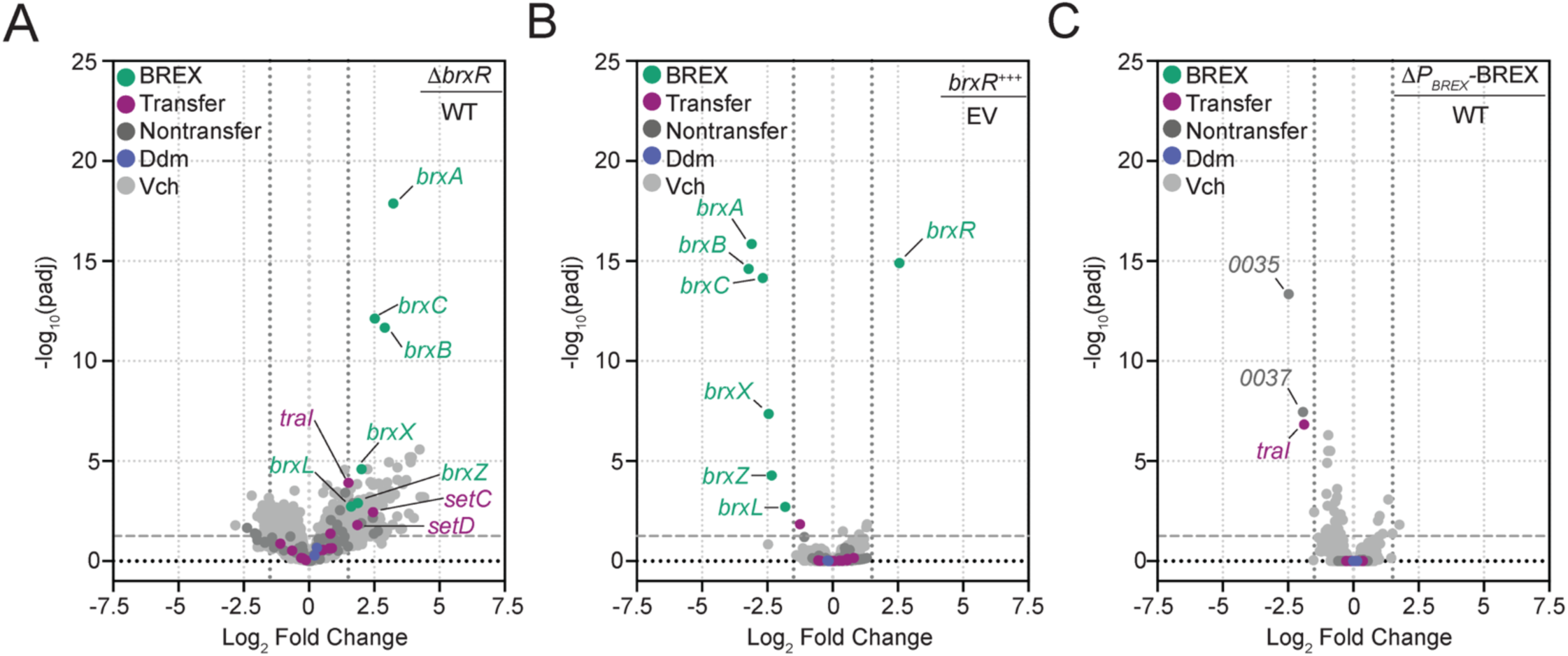
BrxR is responsible for higher BREX expression specifically during phage infection and for repressing the BREX operon; its deletion indirectly alters the host transcriptome. A-C) Volcano plots showing changes in gene expression when comparing **A)** the Δ*brxR* mutant to the wild-type strain (WT), **B)** a strain expressing BrxR *in trans* (*brxR^+++^*) to a control strain with an empty expression cassette (EV), and **C)** the Δ*P*_BREX_-BREX strain (lacking the BREX promoter region and BREX system) to the wild-type strain (WT). Vch denotes genes encoded in the *V. cholerae* chromosome outside of the SXT ICE. −Log_10_ (p_adj_) ≥ 1.25 and Log_2_ fold change ≥ ± 1.5 were considered significant (grey dashed and dotted lines, respectively). Differential expression was determined using DESeq2^29^. See Supplementary Tables 8, 10, and 11 for exact values.

Because BrxR overexpression specifically repressed the BREX operon, we hypothesized that the widespread transcriptional changes in the Δ*brxR* mutant were an indirect consequence of elevated BREX abundance. To test this, we compared the transcriptomes of wild-type cells and a mutant lacking BrxR and the entire BREX system, including its promoter (Δ*P_BREX_-BREX*; Fig. 4C). In sharp contrast to the >200 genes misregulated in the Δ*brxR* mutant (Fig. 4B), deletion of BREX along with *brxR* restored gene expression to near wild-type levels, with only five genes differentially expressed. These findings demonstrate that the global transcriptional changes seen in the absence of BrxR are due to derepression and accumulation of BREX. Overall, our results show that BrxR is a dedicated repressor of the BREX system and does not directly regulate other host genes.

We next tested whether these transcriptomic perturbations or unregulated BREX levels translate to a measurable host fitness cost. In competition assays between wild-type and Δ*brxR* mutants, we detected no fitness cost from constitutively high BREX expression in either rich or defined media (Extended Data Fig. 5C and D). Thus, despite widespread transcriptional changes, increased BREX abundance is not inherently detrimental to host fitness under these conditions.

Since phage defense systems are commonly encoded on MGEs^23,46,47^, we next asked whether BrxR regulation is important during horizontal transfer into naïve hosts, where unregulated BREX could, in theory, cause autoimmunity before protective methylation is established^21^. Conjugation frequencies for SXT ICE *Vch*Ind5 encoding wild-type versus unregulated BREX (Δ*brxR*) were indistinguishable (Extended Data Fig. 5E). To rule out the possibility that elevated expression of SXT transfer activators (*setC* and *setD*^36,37^) in the Δ*brxR* donor (Fig. 4B) masked potential toxicity through hyper-transfer, we complemented the Δ*brxR* SXT ICE donor with an ectopic copy of *brxR*, which yielded the same conjugation frequency (Extended Data Fig. 5E). Similarly, pre-expressing BrxR in recipient cells to repress incoming BREX provided only a minor increase in acquisition of the Δ*brxR* element (Extended Data Fig. 5F). Together, these findings demonstrate that BrxR-mediated repression is not required to prevent severe costs of constitutive defense expression. By ruling out baseline toxicity, metabolic burden, and autoimmunity during horizontal transfer, our data support a model in which BrxR’s primary function is to enable dynamic immune priming during restricted phage infection.

### Independent defense-associated WYL regulators coordinate multi-layered defense responses across bacterial genomes

Although BrxR functions as a dedicated repressor of the BREX operon, restricted phage infection also led to the co-induction of the anti-plasmid defense system DdmDE^30^ (Fig. 1A). Because *ddmDE* is not regulated by BrxR (Fig. 4), we hypothesized that *ddmDE* is independently controlled by its own upstream WYL protein (WYL^DE^) in response to the same restricted infection signal. To test this hypothesis, we modeled the structure of WYL^DE^ bound to ssDNA and compared it with our solved structure of ssDNA-bound BrxR (Fig. 5A). Despite sharing only 25.2% amino acid identity, the predicted ssDNA-bound conformation of WYL^DE^ is structurally homologous to BrxR, with the ssDNA electrostatic map from BrxR fitting cleanly into the predicted WYL^DE^ binding pocket. To assess whether WYL^DE^ regulates *ddmDE* through ssDNA interaction, we introduced mutations into the predicted ssDNA-binding pocket of WYL^DE^ (WYL^DE,mut^) analogous to those in BrxR^mut^ (Fig. 5A). Transcriptional profiling during restricted phage infection revealed that WYL^DE,mut^ abolished *ddmDE* induction, whereas BREX induction remained fully intact (Fig. 5B). These results demonstrate that independent WYL repressors sense the same restricted infection through a shared molecular cue (involving ssDNA) to coordinate a heightened immune response across functionally distinct defense systems without direct regulatory cross-talk.

**Figure 5.**
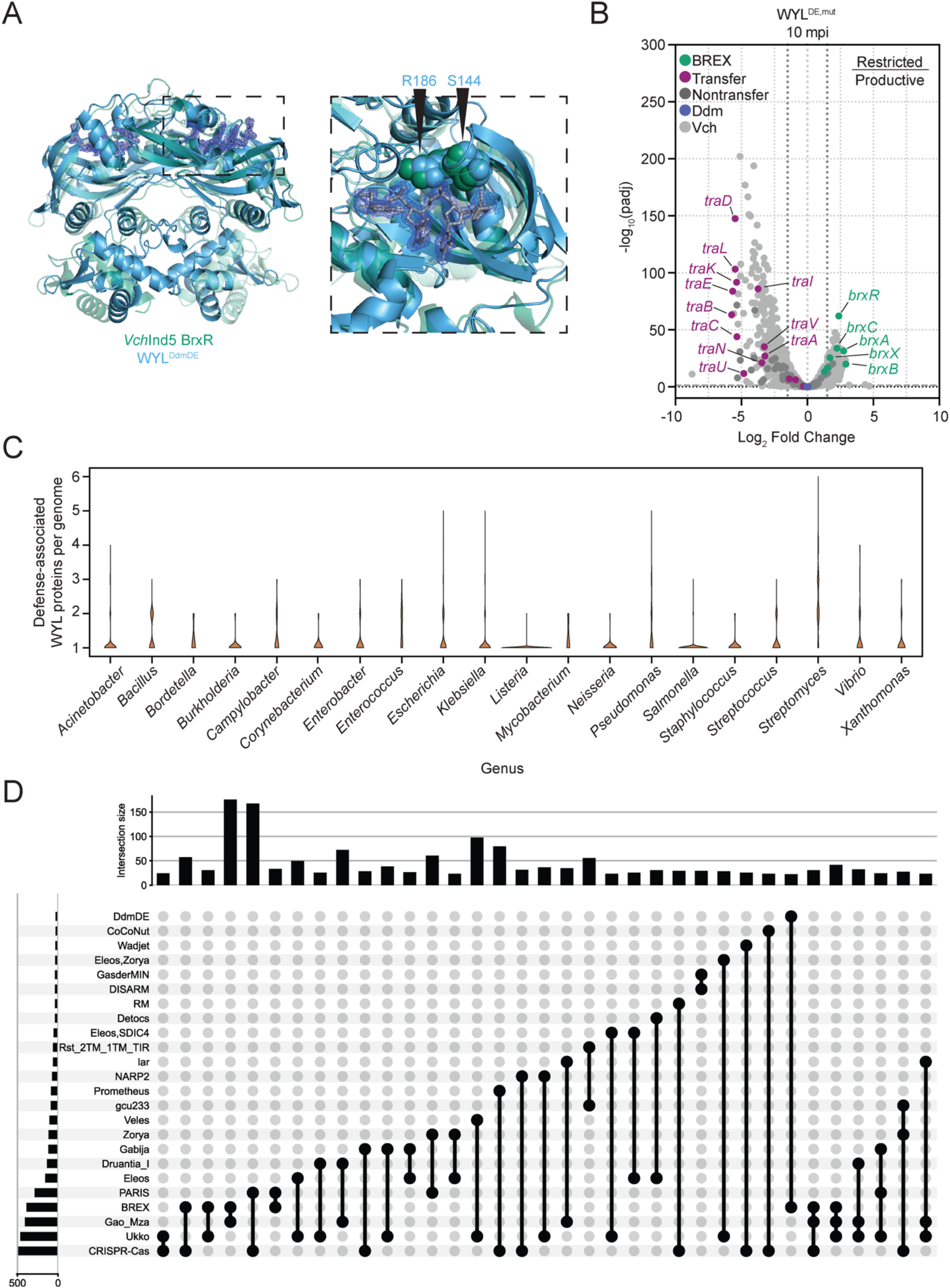
Independent WYL proteins can coordinate the expression of distinct defense systems during a restricted infection. **A)** Predicted structure of the DdmDE WYL (WYL^DE^) bound to ssDNA using AlphaFold 3 (ipTM = 0.83, pTM = 0.85) overlayed on the solved structure of the ssDNA-bound BrxR. The inlet is a zoom-in on the ssDNA binding pocket, with the electrostatic map of the ssDNA from the BrxR structure displayed, and the analogous residues in WYL^DE^ that were mutated in the ssDNA-binding mutant (WYL^DE,mut^). The ssDNA from the predicted structure of WYL^DE^ is not shown. **B)** Volcano plots showing changes in gene expression in the WYL^DE,mut^ background, plotted as the Log_2_ fold change of the restricted (ICP1 Δ*orbA*) relative to the productive (ICP1) phage infection, at 10-minute post-infection (mpi). Differential expression was determined using DESeq2^29^. SXT transfer (*tra*) genes were assigned based on the following references^24,32,33^. Genes encoded by SXT that are not involved in SXT transfer or BREX are categorized as nontransfer. Vch denotes genes encoded in the *V. cholerae* chromosome outside of the SXT ICE. −Log_10_ (p_adj_) ≥ 1.25 and Log_2_ fold change ≥ ± 1.5 were considered significant (grey dashed and dotted lines, respectively). See Supplementary Table 12 for the exact values. **C)** Violin plot showing the distribution of the number of defense-associated WYL proteins per genome for the twenty genera with the highest total representation of these proteins (see Supplementary Table 13 for exact values). **D)** UpSet plot showing the co-occurrence of defense systems that are associated with distinct daWYL proteins in the same genome. Defense systems that flank the same daWYL protein are listed as defense 1 and defense 2 (e.g., Eleos, Zorya). Intersection size indicates the number of genomes that represent a given combination. For the full list of defense combinations per genome, see Supplementary Table 14.

Because *ddmDE* is part of the core defense repertoire in seventh pandemic El Tor (7PET) *V. cholerae*^31^, and SXT/R391 ICEs are widespread in 7PET strains^48^, this coordinated multi-system response appears to be a common feature of epidemic *V. cholerae*. However, whether encoding multiple defense-associated WYL (daWYL) proteins within a single genome is a broader phenomenon among bacteria remained unknown. To evaluate the distribution of daWYL regulators, we identified all WYL proteins across complete bacterial genomes in NCBI RefSeq^49^, and filtered for those encoded immediately adjacent to a defense system, inferring that these proteins likely regulate the neighboring defense. This stringent approach defined a set of 13,508 (Supplementary Table 13) unique putative daWYL proteins. Assessing their genomic distribution revealed that daWYL-encoding genomes typically encode 1 to 5 distinct daWYL proteins, reaching a maximum of 11 in a single *Paenibacillus* genome (Fig. 5C and Supplementary Table 14). Notably, each of the twenty most well-represented genera included genomes encoding two or more daWYL proteins, indicating that coordinated WYL-mediated defense regulation extends to diverse taxa.

To map the defense repertoire potentially governed by these daWYL regulators, we analyzed the co-occurrence of distinct defense systems paired with independent daWYL proteins (Fig. 5D and Extended Data Fig. 6A). Most co-occurrences involved pairings of two defense systems, consistent with the overall genomic distribution (Fig. 5C). Among combinations present in 20 or more genomes, the most frequent pairings were BREX with Gao_Mza^2^, and CRISPR-Cas with PARIS^50^ (Fig. 5D and Supplementary Table 15). To identify less abundant co-occurrences, we examined combinations present in 5 to 20 genomes, revealing multiple instances where single genomes encode 4 or 5 distinct defense systems each proximal to a WYL protein (Extended Data Fig. 6A). Finally, to determine how many of these daWYL proteins belong to the most well characterized BrxR and CapW families, we performed pairwise Foldseek^51^ structure-plus-sequence alignments (Extended Data Fig. 6B). Clustering of the resultant pairwise distances yielded 47 distinct clusters. BrxR, CapW, and WYL^DE^ clustered together, representing 12.5% of total daWYL proteins, while several remaining daWYLs formed even larger, uncharacterized clusters. Together, these findings indicate that the coordination of distinct defense systems by independent WYL regulators is a widespread bacterial strategy, driven by diverse and largely uncharacterized families of daWYL proteins.

## Discussion

WYL-domain regulators are associated with a wide range of defense systems across bacterial phyla, with nearly 30% of complete prokaryotic genomes encoding at least one defense-associated WYL protein (Supplementary Table 14). While previous studies have established that WYL-domains bind ssDNA *in vitro*^22,52,53^ and increased immune protein abundance enhances defense in several systems^15,18,22,41,54^, the natural cues that trigger derepression—and the extent to which inducible immunity provides a selective advantage in native host-phage contexts—have remained unclear. Here, we demonstrate that restricted phage infection serves as a natural, physiologically relevant trigger for WYL-mediated defense derepression. In its endogenous genetic context, this dynamic response elevates defense protein abundance, shifting host-viral stoichiometry and priming surviving bacteria to defeat phages encoding anti-defense proteins such as OrbA (Fig. 6). By exploiting a primary restricted infection as a clear signal of a phage-dense environment, bacteria can dynamically amplify immunity precisely when evolved phages with counterdefense mechanisms are most likely to attack.

**Figure 6.**
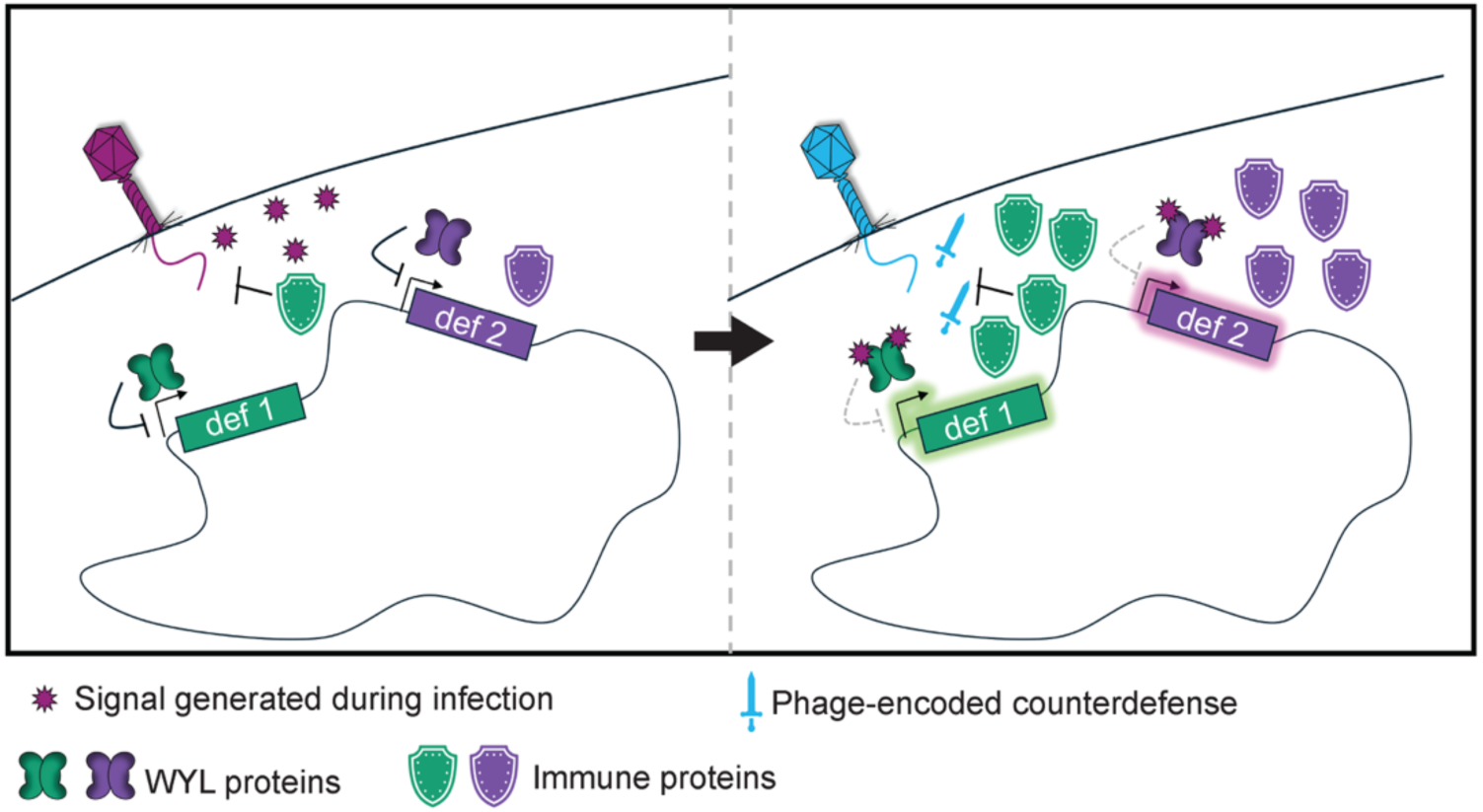
Primed immunity induced by a restricted infection provides heightened protection against phage-encoded inhibitors. Model of a coordinated induction of defense systems during a restricted infection by independent WYL proteins. In the absence of phage infection, WYL repressors maintain their cognate defense system at a basal expression level. During a restricted infection (e.g., an inhibitor-lacking phage; magenta), a signal is generated (starburst) that can be sensed by multiple, distinct WYL repressors. This sensing inhibits the repressor’s activity, leading to the derepression of their cognate defense system (e.g., ‘Defense 1’ and ‘Defense 2’) and establishing a heightened, multi-layered immune state. Upon a subsequent infection by an inhibitor-proficient phage (blue), the heightened immune state bolsters protection against the phage’s counterdefense, thereby restricting the infection and ensuring cell survival. Def −Defense

Our findings reveal an expanded regulatory framework that allows defense systems to coordinate a broader immune response when phage infection is imminent. Because restricted infection produces a shared molecular signal, independent WYL repressors can sense this cue in parallel, without regulatory crosstalk. This modular architecture clarifies the biological utility of WYL-regulated abortive infection systems such as CBASS. When an abortive system is the sole frontline defense, increased protein abundance offers no benefit as the cell is destined to die following primary infection. However, when a non-lethal frontline defense like BREX clears the infection and the host survives, bystander WYL repressors harness the shared signal to upregulate their associated systems, preparing the surviving host to fend off subsequent, distinct phage attacks (Fig. 6). It remains to be determined whether the specific mechanism of primary restriction influences cue generation and downstream WYL activation, potentially tuning this systemic response.

Diverse MGEs disseminate defense systems in bacteria^23,46,47^, and our findings shed light on how these elements may respond to different infection outcomes. Notably, productive and restricted phage infections trigger divergent regulatory programs, with productive infection leading to downregulation of BREX and strong induction of genes necessary for horizontal transfer. The specificity of BREX induction to restricted infection is surprising, given that productive infection, where ICP1 degrades the bacterial genome^26^, should also generate the known ssDNA ligand for WYL proteins such as BrxR. This is further complicated by robust activation of SXT transfer genes during productive infection without detectable upregulation of their canonical master activators, SetC and SetD. While the canonical SXT transfer pathway relies on the host SOS response to alleviate repression of *setCD*^37^ and to activate conjugative transfer^33^, our data suggest an alternative, phage-specific response. A compelling hypothesis is that SXT ICEs, and possibly other defense-associated MGEs, have evolved a regulatory hierarchy to ensure that when the host is defensible, its immunity is amplified, but when infection is productive and host death is inevitable, the element suppresses futile defense expression and prioritizes horizontal transfer. This could explain both the induction of transfer genes and the suppression of the BrxR-mediated defense, even if BrxR’s activating signal is present. Such a mechanism would allow MGEs to distinguish between distinct lethal threats, such as DNA damage versus viral takeover, and tailor their responses accordingly.

Beyond immune priming, our findings challenge the prevailing assumption that transcriptional repression of bacterial defenses evolved mainly to mitigate host toxicity. Previous studies relying on heterologous expression platforms have reported severe growth defects or cell death when repressors for systems such as CBASS, CRISPR-Cas, and other BREX systems are inactivated^15,16,18^. In contrast, we found that constitutive, high-level BREX expression from its native ICE locus incurred no detectable fitness cost. This suggests that observed toxicities in non-native contexts may result from artificial expression systems or imbalanced stoichiometries rather than an inherent cost of defense expression. Thus, toxicity mitigation is unlikely the primary evolutionary driver of BREX repression. Instead, our work points to both host-centric and evolutionary rationales for dynamic regulation. While constitutive expression of a single defense system can be well tolerated, we hypothesize that sustained, unregulated co-expression of multiple active defense complexes across the genome, such as following restricted infection, could impose synergistic fitness costs or increase the risk of autoimmunity. From an evolutionary perspective, constitutive, high-level defense expression would subject phage populations to continuous selective pressure, accelerating the evolution of more effective counterdefenses, such as inhibitors with higher binding affinity or elevated expression levels. WYL-mediated dynamic regulation alleviates this risk by confining broad immune activation to periods of active viral threat, thereby minimizing the selective pressure for counterdefense evolution while preserving the host’s capacity to mount a potent defense response in phage-rich environments.

## Materials and methods

### Bacterial strains and growth conditions

All *V. cholerae* and *E. coli* strains were either grown in Lysogeny Broth (LB) with aeration or on LB plates supplemented with 1.5% agar (Fisher BioReagents). *V. cholerae* strains are derived from strain E7946. Antibiotics, when used, were added at the following concentrations: 100 μg/mL streptomycin, 75 μg/ml Kanamycin, 100 μg/mL spectinomycin, 32 μg/mL trimethoprim. When an inducer was required for the P*_tac_-riboE* promoter, either 1 mM β-D-thiogalactopyranoside (IPTG) with 1.5 mM theophylline or, when stated, a 10-fold dilution of both inducers was used. For the P*_bad_-riboE* promoter, 0.2% arabinose plus 1.5 mM theophylline was used. Cells were induced at the time of back dilution and maintained throughout all other growth conditions. The genotypes of all strains used in this study are included in Supplementary Table 8.

### Plaque assays

Phages used in this study are listed in Supplementary Table 8. Phage stocks were stored in STE buffer (100 mM NaCl, 10 mM Tris-Cl pH 8.0, and 1 mM EDTA) at 4°C. Plaque assays used to calculate the efficiency of plating were performed by growing either *V. cholerae* or *E. coli* to an OD 0.3 – 0.5, mixing 50 μL of cells with 10 μL of serially diluted phage in LB, incubated at room temperature for ∼ 10 min to allow for phage adsorption, and then each dilution was added to molten 0.5% top agar (0.7% top agar was used for the *E. coli* phage T7) in a single petri plate. Plaque assays were incubated at 37°C (or at room temperature for the phage T7) overnight and then enumerated the following day. EOP was calculated as a ratio of the number of plaques on the experimental conditions over the number of plaques calculated for the permissive condition (E7946 for *V. cholerae* and MG1655 for *E. coli*). When necessary, 1 mM IPTG and 1.5 mM theophylline, or a 10-fold dilution as stated in the figure legend, were added to molten 0.5% top agar to maintain induction conditions.

### Generation of mutant strains

Deletions in the *V. cholerae* chromosome were generated by amplifying 1 kb arms of homology by PCR and a spectinomycin antibiotic resistance cassette flanked by FRT recognition sites (spec cassette). Introduction of affinity tags and point mutations at native sites was achieved by amplifying 2kb arms of homology, inserting either the affinity tag DNA or the point mutation sequence in the middle of the two homology arms. For the deletion, the arms of homology and the spec cassette were stitched together using splicing by overlap extension (SOE) PCR. For the native site constructs, the homology arms were stitched together using SOE PCR. Both constructs were then introduced into *V. cholerae* by natural transformation^55^. The native site constructs were co-transformed with a selectable marker (*lacZ::spec* or *lacZ::kan*), where the native construct was present at a 3- to 5-fold excess over the selectable marker. *V. cholerae* cells were made naturally competent by growing the cells in LB at 30 °C until an OD ∼0.3 – 1.0, pelleting the cells at 5000 x *g* for 3 minutes, washing the cells in 0.5X Instant Ocean, then adding the cells to chitin (Sigma) slurry in 0.5X Instant Ocean and incubating for 24 hours at 30 °C. Transformants were selected by plating an LB agar plate with the appropriate antibiotic. Native site constructs were screened by colony PCR; colonies that exhibited the proper banding pattern were then Sanger sequenced to verify that the tag or corresponding mutation was present and that no mutations had been introduced in the homology arms.

Identification of the BREX promoter was done using transcriptomic data to approximate the start of the BREX operon (see RNA sequencing analysis for details). The putative promoter region was identified using SAPPHIRE^56^, the identified promoter region along with the entire BREX system was deleted using the deletion protocol above.

### RNA sample collection, isolation, and sequencing

*V. cholerae* strains were grown to an OD of 0.3 and infected with either ICP1 or ICP1 Δ*orbA* at a multiplicity of infection of 2.5 and incubated at 37 °C at 220 rpm. Five mL of the sample was removed at the time of addition of phage (serving as the zero-minute time point), mixed with equal volumes of ice-cold methanol, and pelleted at 5000 x *g* for 5 minutes at 4 °C. The supernatant was removed, and the pellets were resuspended in 1 mL of ice-cold 1x phosphate-buffered saline (pH 7.4). Samples were then pelleted again at 5000 × *g* for 5 minutes at 4 °C. The supernatant was removed, and the pellets were resuspended in 200 μL of TRI Reagent (Sigma/Millipore). The samples were then stored at −80 °C. Samples were thawed for 10 minutes at room temperature, then 40 μL of chloroform was added, vortexed and incubated at room temperature for an additional 10 minutes. Samples were centrifuged at 12,000 × *g* for 10 minutes at 4 °C. From the phase-separated samples, the top aqueous layer was mixed with 110 μL of isopropanol and 11 μL of 3 M sodium acetate (pH 6.2) and then mixed by vortexing. Samples were centrifuged at 12,000 × *g* for 15 minutes at 4 °C, and the RNA pellets were washed twice with 75% ethanol. Excess ethanol was removed from RNA pellets by incubating at 65 °C for 3 minutes. RNA was rehydrated in 20 μL of Diethyl dicarbonate (DEPC)-treated nuclease-free water. The concentration and integrity of the purified RNA were measured using a Nanodrop. For mitomycin C (MMC) treatment, cells were grown to an OD of 0.3, and an initial 5 mL sample was removed. MMC was added to the culture to a final concentration of 20 ng/mL, and the culture was incubated at 37°C with aeration for 60 minutes; then a 5 mL sample was removed. RNA was isolated following the procedure above. Sequencing was performed by SeqCenter (Pittsburgh, PA) at a sequencing depth of 12 million reads. Raw data were demultiplexed, quality-checked, and adapters trimmed by SeqCenter.

### RNA sequencing analysis

The paired-end FASTQ read files were mapped to the reference sequences for the *V. cholerae* chromosomes 1 (CP024162.1) and 2 (CP024163.1), and the integrative conjugative element SXT *Vch*Ind5 (GQ463142) using bowtie2^57^. The resulting mapped alignment files (.bam) were sorted using samtools^58^, and then the estimation of transcript abundance was determined by StringTie^59^. The transcript count matrix was generated using the associated StringTie Python script prepDE.py, which generated an estimated transcript count matrix. The transcript count matrices were then imported into RStudio for analysis using the DESeq2^29^ package.

Differential sequencing analysis was performed using the DESeq2 package. For phage infections, the respective time points for ICP1 and ICP1 Δ*orbA* were compared to each other (e.g., ICP1 4 minutes post-infection vs. ICP1 Δ*orbA* 4 minutes post-infection). Genes that were considered to be differentially expressed had a Log_2_ fold change of less than or equal to −1.5 or greater than or equal to 1.5, and a −Log_10_ (p_adj_) value greater than 1.25. Volcano plots were generated using GraphPad Version 10.6.0 (769). Heatmaps were generated by applying a variance-stabilizing transformation to transcript count matrices and then Z-score normalized using Pheatmap. The viridis color palette was used to color the heatmaps.

Read tracks were generated by indexing and sorting .bam files using samtools, and scaling factors were estimated using BEDtools^60^. The forward and reverse-strand bedgraph files were generated with BEDtools and the determined scaling factors. The averaged bedgraph files were then visualized in JBrowse 2^61^.

Gene ontology (GO) enrichment analysis was performed in RStudio using the clusterProfiler Package^62^. Functional annotations of the *V. cholerae* E7946 and SXT *Vch*Ind5 proteome were generated by submitting the pooled proteome to eggNOG-mapper v2.1.13^63^ using DIAMOND^64^ for sequence similarity searches against the eggNOG v5.0 database^65^ at the Vibrionales taxonomic level. Additionally, GO terms mapped from the *V. cholerae* N16961 proteome (Uniprot ID 243277) via ortholog correspondence were also used. Over-representation analysis was performed separately for upregulated and downregulated differentially expressed genes with p_adj_ < 0.05 and Log2 Fold Change > 1, using the Benjamini-Hochberg method to control the false discovery rate. GO term descriptions were retrieved using the GO.db Bioconductor package.

### Western blot and densitometry analysis

*V. cholerae* cells were grown to an OD of 0.3 as stated above in growth conditions. During phage infections, an MOI = 5 was used. At each time point, 5 mL of the sample was spun down at 5000 × *g* for 3 minutes. Pellets were resuspended in 20 μL of lysis buffer (50mM Tris-HCl, pH 7.6, 150 mM NaCl, 1 mM EDTA, pH 8.0, 0.5% Triton X-100, and a Pierce protease inhibitor EDTA-free tablet (Thermo Fisher Scientific)), and incubated at 65 °C for 10 minutes. Protein concentration was determined by Pierce BCA assay (Thermo Fisher Scientific). Protein samples were diluted to a final concentration of 1 μg/μL using lysis buffer with the addition of 4X Laemmli Buffer supplemented with 2-mercaptoethanol (10%) (final concentration of 1X; Bio-Rad), and boiled at 99 °C for 10 minutes. From each sample, three Any-Kd Mini-PROTEAN TGX Precast gel (Bio-Rad) were loaded with 10 μL of sample (10 μg of total protein) and were run in parallel. Protein gels were then transferred to a Mini-size PVDF membrane (Bio-Rad) using a Trans-Blot Turbo system (Bio-Rad). Proteins were detected using a primary α-Flag antibody (Sigma) at a 1:10,000 dilution, α-HA antibody (Invitrogen) at a 1:10,000 dilution, or a custom polyclonal antibody against the BrxR protein from SXT *Vch*Ind5 (Genscript) at a 1:1000 dilution, and a secondary goat α-rabbit-HRP antibody (Bio-Rad) at a 1:10000 dilution. Blots were developed using Clarity Western ECL substrate (Bio-rad) and imaged using a Chemidoc XRS imaging system (Bio-Rad).

Densitometry analysis was performed using ImageJ and conducted between bands of the same gel, as well as across gels that were run in parallel. Each experiment had its own wild-type control for comparison. For the phage infection time courses, the band densities were measured for the non-specific band recognized by the BrxR antibody, BrxR, BrxC-3xFLAG, and BrxZ-HA, and divided by the zero-minute time for the same protein. Then the ratios for BrxR, BrxC-3xFLAG, and BrxZ-HA were normalized by dividing by the non-specific band ratio. For comparing the BrxR mutants or the BREX overexpression densitometry analysis, the bands were compared to the wild-type band density for each protein, then normalized by the non-specific band ratio. The presented densitometry fold change is the average of three biological replicates.

### Burst assay

*V. cholerae* strains were grown to an OD of 0.3 at 37 °C, then infected with ICP1 at an MOI = 0.1. For subsequent infection, cells were first infected with ICP1 Δ*orbA* or ICP1 Δ*gp15-36* at an MOI = 5 and incubated for 40 minutes at 37 °C, then infected with ICP1 at an MOI = 0.1. Infected cells were incubated at 37 °C for 7 min to allow adsorption. Infected cells were then diluted 1:2500, 1:25000, and 1:250000 in pre-warmed LB at 37 °C. To quantify the number of input phages, 500 μL of the 1:2500 dilution was taken out immediately to perform a plaque assay with the permissive background E7946. After 40 min of incubation at 37 °C, the number of phage progeny produced was measured by removing 500 μL of each dilution and then treating 20 μL with chloroform. After 10 minutes, the cell debris and chloroform were removed by centrifugation at 5000 × *g* for 15 min. Supernatant was removed and used to perform a plaque assay with the permissive background E7946. The number of phage progeny produced was calculated by dividing the number of output phages produced per mL by the number of input phages.

### Phage genome replication after subsequent infections by quantitative polymerase chain reaction

*V. cholerae* strains were grown to OD 0.3 and infected with either ICP1 Δ*orbA* or ICP1 Δ*gp15-36* at a multiplicity of infection of 5 and vortexed. Infected cultures were incubated at 37°C with aeration for 40 minutes. Cells would then be infected with ICP1 at a multiplicity of infection of 0.1 and vortexed. Immediately, a 100 μL sample was removed and served as the zero-minute time point and boiled at 95°C for 10 minutes. Cultures were incubated at 37°C with aeration, and 100 μL samples were removed 20 minutes post-secondary infection, then boiled at 95°C for 10 minutes. A standard of purified ICP1 genome was made by 10-fold dilutions, while boiled samples were diluted 50-fold to serve as a template for qPCR. ICP1 genome replication was measured with *orbA* specific primers RO822(CTGTGACAGTCTACAATCATGG)/823 (CTCGACTCATCCCTTCATCA) using iQ SYBR Green Supermix (Bio-Rad) and the CFX Connect RealTime PCR Detection System (Bio-Rad). Two technical duplicates were used for each biological replicate. Fold change in genome copy number was determined by dividing the calculated sq value of each time point by the zero-minute time point.

### Identification of phage mutant with candidate superinfection exclusion (SIE) proteins deleted

To identify candidate genes that encode a SIE protein, we compared ICP1 gene expression during restricted and productive infections for our dataset (Supplementary Tables 1 and 2) and a recent preprint^66^. From this analysis, we identified 13 ICP1 genes that were not differentially expressed. Moreover, these genes were located in a genomic region near where the BREX-inhibitor OrbA is encoded, a protein that is expressed early in infection. From our collection of ICP1 deletions, we identified a phage with a deletion spanning most of the genes that were not differentially expressed (Δ*gp15-36*), including *orbA* (*gp25*).

### SXT conjugation assay

SXT conjugation assays were performed as previously described^21,37^. SXT transfer between *V. cholerae* and *V. cholerae* is low^21,37^, so to increase the conjugation frequency, we incubated the donor cells with mitomycin C before mating (MMC). Specifically, overnight cultures of donor strains were incubated at 37 °C with MMC at a final concentration of 20 ng/μL and then incubated at 37 °C for 15 minutes. 500 µL of donor cells were first pelleted and washed with fresh LB at 5000 × *g* for 3 minutes at room temperature. Then, they were mixed with an equal volume of the recipient cells, which had been grown overnight at 37 °C. Cells were pelleted at 5000 × *g* for 3 minutes at room temperature. The supernatant was removed, and the cells were resuspended in 50 μL of fresh LB. They were then spread onto a 0.8 μm filter disc placed on top of an LB plate. Matings were incubated at 37 °C for 6 hours. The filter disc was then placed in 1 mL of fresh LB, vortexed, and then different volumes were plated on LB agar plates supplemented with trimethoprim and either spectinomycin or kanamycin. In cases where induction was required, the inducer was included in the LB agar plates.

### Competition assay

*V. cholerae* strains were grown overnight in liquid LB at 37 °C. Strains were back-diluted to an OD of 0.005 into the same flask with fresh LB. Cells were grown at 37 °C, shaking at 220 rpm. At the indicated time points, 200 μL of the sample was removed, serially diluted, and plated on LB agar plates supplemented with X-gal (40 μg/mL), streptomycin, and trimethoprim. Plates were incubated overnight at 37 °C. *V. cholerae* strains that harbored a wild-type SXT encoded a functional copy of LacZ, and colonies appeared blue. *V. cholerae* strains that harbored an SXT where WYL is absent appeared white as *lacZ* was disrupted with an antibiotic resistance cassette. For competition assays in M9 minimal media (1X M9 salts (Sigma Aldrich), 1X MEM vitamin solution (Gibco by Life Technologies), 2mM MgSO_4_, 0.1mM CaCl_2_, 0.4% glucose)) 1 mL from an overnight culture grown in LB was washed twice with 2 mL of M9 minimal media to remove excess nutrients at 5000 x g for 3 minutes. The cell pellets were resuspended in 1 mL of M9 minimal medium and then back-diluted to an OD of 0.005 in the same flask with fresh M9 minimal medium. The competitive index was calculated by first dividing the number of colony-forming units (CFU) per mL calculated for the WYL-mutant background by the CFU/mL for the wild-type background at each time point, then dividing the ratio of each time by the ratio of the two strains at the zero-hour time point.

### Protein expression and purification

Codon-optimized BrxR^Ind5^ (Genscript) was subcloned into the pET15b expression vector to contain an N-terminal, thrombin-cleavable 6XHis tag. The BrxR^Ind5^(K197A/S156E) double mutant was generated in this construct by site-directed mutagenesis using NEB’s Q5 mutagenesis kit. Protein expression in *E. coli* BL21(DE3) pLysS cells and purification were carried out as described^17^. Briefly, proteins were purified by metal affinity chromatography followed by thrombin cleavage to remove the 6×His tag. The cleaved protein was further purified by size-exclusion chromatography (SEC) on a HiLoad 16/60 Superdex 200 prep grade column (Cytiva) equilibrated in 25 mM Tris (pH 7.5) and 200 mM NaCl. Peak fractions were pooled and concentrated in an Amicon Ultra centrifugal filter (10,000 molecular weight cutoff; Millipore). Single-use aliquots were flash frozen in liquid nitrogen and stored at −80°C.

### DNA Binding Assays

Electrophoretic Mobility Shift Assays (EMSA, or gel shifts) were used to assess DNA binding by BrxR^Ind5^. A 29 bp sequence corresponding to the predicted BrxR^Ind5^ target site, located upstream of the *BrxR^Ind5^*‘s coding sequence, was embedded within a 60 bp dsDNA oligo (acgatgcagaacagttcTGATACCTAATAAATTTAATTAGGTTGAAtaaatgatgtccat). Additional non-specific DNA was included on either side of the target sequence to minimize end effects on DNA binding. A non-specific “scrambled” oligo containing the same flanking sequence was also tested in binding assays (acgatgcagaacagtGATTGGCTTATAAATTCTGGCATAACGGGAtaaatgatgtccata). dsDNA substrates were generated by annealing complementary ssDNA oligonucleotides (IDT Inc.) resuspended in 10 mM Tris, 0.1 mM EDTA (pH 8.0). Equal amounts of each oligonucleotide (500 pmol; 5 µL of a 100 µM stock) were combined in a final volume of 50 µL with nuclease-free water. Annealing was performed in a thermal cycler by heating samples to 95°C for 5 min, followed by gradual cooling at a rate of 0.5°C every 30 s for 170 cycles, with a final hold at 25°C. Annealed dsDNA was purified using a DNA Clean & Concentrator kit (Zymo Research) and eluted in 50 µL of 10 mM Tris (pH 8.5). DNA concentration was quantified using a Qubit dsDNA High Sensitivity assay kit (Invitrogen) and samples were normalized to 500 nM. Purity and annealing quality were assessed using an 8% native polyacrylamide gel.

For DNA-binding reactions, dsDNA substrates were used at a final concentration of 50 nM and incubated with purified BrxR^Ind5^ at concentrations ranging from 12.5 nM to 1 µM (diluted in 100 mM NaCl, 25 mM Tris, pH 7.5) in binding buffer containing 20 mM Tris (pH 7.9), 40 mM NaCl, and 2.5% sucrose. Samples were incubated at 23°C for 30 minutes prior to separation on 8% native acrylamide gels (29:1 acrylamide:bisacrylamide; Bio-Rad) prepared in 0.5 X TBE.

For reactions testing the effect of ssDNA on BrxR^Ind5^ binding to its dsDNA target, 800 nM BrxR^Ind5^(WT) or 1000 nM The BrxR^Ind5^(K197A/S156E) was incubated with 50 nM dsDNA target sequence and a 7-mer poly-T ssDNA (concentrations ranging from 500 µM to 25 nM) for 30 minutes prior to loading samples onto gels.

Gels were pre-run for 15 minutes at 70 V in 1 X TBE, after which wells were rinsed prior to sample loading. 5 µl of each binding reaction was loaded directly on the gel without loading dye and electrophoresed at room temperature for 60 min at 70 V in 1× TBE buffer. DNA was visualized by staining gels with SYBR Gold (Invitrogen) diluted in 1× TBE for 30 min, followed by two rinses in 1× TBE and imaging using a Bio-Rad gel documentation system.

EMSA reactions were quantified by densitometric analysis in ImageJ. The fraction of DNA bound by BrxR^Ind5^ was determined using the equation Y = 1 - (IT/IC), where IT represents the intensity of the unbound probe band in experimental lanes and IC corresponds to the intensity of the unbound probe in the no-protein control lanes^16^.

### Crystallographic Structure Determination

Crystals of apo-BrxR^Ind5^ were grown by hanging drop vapor diffusion over 1 mL well solutions. Drops containing 1 µL of a solution of 10 mg/mL (150 µM) BrxR^Ind5^ and 1.5 mM 3’3’ cyclic-GAMP were mixed with 1 µL of reservoir solution containing 0.1 M malic acid (pH 7.0) and 16.25% (w/v) PEG3350. Coverslips containing crystals were moved to wells with sequentially higher PEG 3350 concentrations (25%, 30%, 35%, 40%), while maintaining a constant malic acid concentration and pH, allowing at least 24 hours between each transfer for equilibration. Crystals were flash frozen in liquid nitrogen directly from their growth drops after a final incubation over 0.1 M malic acid (pH 7.0) and 40% (w/v) PEG 3350.

A 6-mer single-stranded DNA oligonucleotide (5’-TTTTTT-3’) was pre-incubated with 125 µM BrxR^Ind5^ at a 3.6-fold molar excess for 30 minutes, and hanging drops were prepared as described above with a well solution containing 0.1 M malic acid (pH 7.0) and 20% (w/v) PEG 3350. Prior to flash freezing in liquid nitrogen, crystals were transferred to a cryoprotection solution containing 0.1 M malic acid (pH 7.0), 20% PEG 3350 and 22% (w/v) ethylene glycol for 1 minute.

Data were collected at the Advanced Light Source synchrotron facility (Berkeley, CA) on beamline 5.0.1 at a wavelength of 0.97Å. Apo-BrxR^Ind5^ was integrated, scaled and merged using HKL2000^67^. BrxR^Ind5^-ssDNA was auto-processed with XDS (data integration and scaling^68^), POINTLESS (space group determination^69^), and AIMLESS (final merging^70^) with a resolution cutoff of I/sigI >2.0. Data was phased via molecular replacement with Phaser^71^ using an ensemble created from all eight chains within the asymmetric unit of a crystallographic model of the BrxR homolog from *Escherichia fergusonii* (PDB ID 7QFZ)^16^. Iterative rounds of refinement with phenix.refine and rebuilding with Coot were performed^72,73^. Crystallographic data is listed in Supplementary Table 9.

### Alphafold3 and protein structure

The primary sequences of DdmDE WYL protein (WP_001277248.1) from *V. cholerae* E7946 were submitted to the AlphaFold3^74^ and were visualized using Chimera X^75^. A protein structure overlay was made using the structural analysis tool Matchmaker in Chimera X. Root mean square deviation (RMSD) was calculated using the standard Matchmaker parameters.

### Distribution of defense-associated WYL proteins

All complete non-metagenomic prokaryotic genome protein FASTA files and GBFF files with NCBI annotations were downloaded from NCBI RefSeq release 230.0 using the NCBI Datasets command-line tool^49,76^. The profile HMM for the WYL domain (defined by PF13280) was then used for an hmmsearch against the protein FASTA files with an E-value inclusion threshold <= 0.001 using HMMER 3.4^77,78^. All WYL protein hit accession numbers were saved. Then, the GBFF files were used to construct a database of every protein sequence upstream and downstream to each WYL hit (agnostic of strand), with taxon labels extracted from the GBFF. All HMM profiles used in system detection in DefenseFinder 2.0^4^ and CRISPRCasFinder 4.3.2^79^ were then used to conduct an hmmsearch for each amino acid sequence collected within the upstream/downstream genes, with the best hit for each with a p-value at or below 0.01 being recorded as that WYL domain-containing protein’s associated defensive gene(s) (proximal WYL hits excluded)^4,77,79^. The package seaborn in Python 3.12.2 was used to plot the distribution of total WYL hits and defense-associated hits per genome for a chosen subset of genera as a violin plot^80,81^.

For the 13,508 putatively defense system-proximal unique WYL protein amino acid sequences, a 3Di structure database was created using Foldseek^51^ with the ProstT5^82^ sequence-to-structure model. The Foldseek search function with default parameters was used to compute pairwise alignments between each protein. The resulting bitscores between the aligned sequences were used to compute a distance matrix, which was used to produce a t-SNE embedding in two dimensions with the Scikit-learn package in Python 3.12.2 with a perplexity of 200. NetworkX was used to build a graph using the distance matrix with an edge threshold of 0.30, and then the NetworkX implementation of Louvain community detection was used to generate clusters from the resultant graph. The t-SNE embedding with clusters defined by Louvain communities was then plotted using Matplotlib 3.10.9

## Data availability statement

The sequencing data generated for this work can be found in the Sequence Read Archive under BioProject accession PRJNA1354459 (Reviewer link https://dataview.ncbi.nlm.nih.gov/object/PRJNA1354459?reviewer=3h8v8g5hlrsv0dfb86ueuau896). Final structures were deposited and validated by the RCSB PDB^83^ (RCSB.org) with PDB IDs 13DB (apo-BrxR^Ind5^) and 13DC (BrxR^Ind5^-ssDNA).

## Acknowledgments

We thank past and present Seed lab members for insightful discussion and feedback on this project. The project described was supported by Grant number R01AI127652 to K.D.S from the National Institute of Allergy and Infectious Diseases of the National Institutes of Health. Its contents are solely the responsibility of the authors and does not necessarily represent the official views of the National Institutes of Health. This work was supported in part by funding to K.D.S. as a Biohub, San Francisco, Investigator and by the Miller Institute for Basic Research in Science, University of California Berkeley. K.D.S. holds an Investigators in the Pathogenesis of Infectious Disease Award from the Burroughs Wellcome Fund. This work received funding from the National Institutes of Health NRSA Trainee Fellowships (5T32 GM132022 to AC and 5T32 HG000047-25 to AAP). The Berkeley Center for Structural Biology is supported in part by the Howard Hughes Medical Institute. The Advanced Light Source is a Department of Energy Office of Science User Facility under Contract No. DE-AC02-05CH11231. The Pilatus detector on 5.0.1. was funded under NIH grant S10OD021832. The ALS-ENABLE beamlines are supported in part by the National Institutes of Health, National Institute of General Medical Sciences, grant P30 GM124169.

## Author Contributions

Conceptualization: RTO and KDS. Methodology: RTO, AAP, DTD, LAD, CG, and AC. Software: RTO, AAP, DTD, and CG. Investigation: RTO, AAP, DTD, CG, LAD, AC, KMH, JPA, SFM, BKK and SN. Writing – original draft: RTO and KDS. Writing – review & editing: all authors. Visualization: RTO, AAP, and KDS. Supervision: KDS. Funding acquisition: BKK and KDS.

## Notes

### Competing Interest Statement

The authors have declared no competing interest.

### Summary of Updates

The manuscript was significantly revised to include new data, streamline the narrative and remove data regarding counterdefense discovery. Specifically, Figure 2 was updated to include the structural mechanism of WYL derepression, showing that ssDNA binding induces an allosteric shift in BrxR that releases dsDNA binding to drive immune gene expression. Figure 2 includes new data showing that defense priming requires a specific phage-derived cue rather than generic DNA damage, distinguishing this immune response from the host SOS pathway. Figure 5 was replaced with new data showing DdmDE derepression during restricted infection depends on its cognate WYL regulator and new bioinformatic analyses showing that multi-WYL regulation is a widespread bacterial paradigm, in which distinct defense-associated WYL regulators independently coordinate multi-layered immune responses without regulatory crosstalk.

