## Extended data Figures 1-6 and Supplementary Figures 1-5 for "Surviving phage attack primes bacterial immunity to defeat counterdefenses"

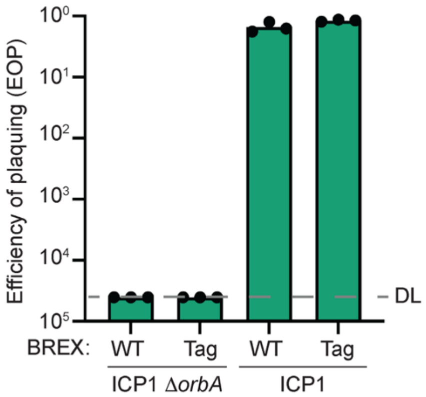


**Extended Data Figure 1. Affinity-tagged BREX functions like untagged BREX system.** Efficiency of plaquing (EOP) of ICP1 and ICP1 Δ*orbA* on the BREX$^{Tag}$ strain (Tag), compared to the untagged BREX system (WT). The bars represent the mean of three independent biological replicates, which are represented by each individual dot. DL – detection limit. A Student's t-test was performed comparing the BREX$^{Tag}$ strain to the wild type, and the results were not significant ($P > 0.05$).

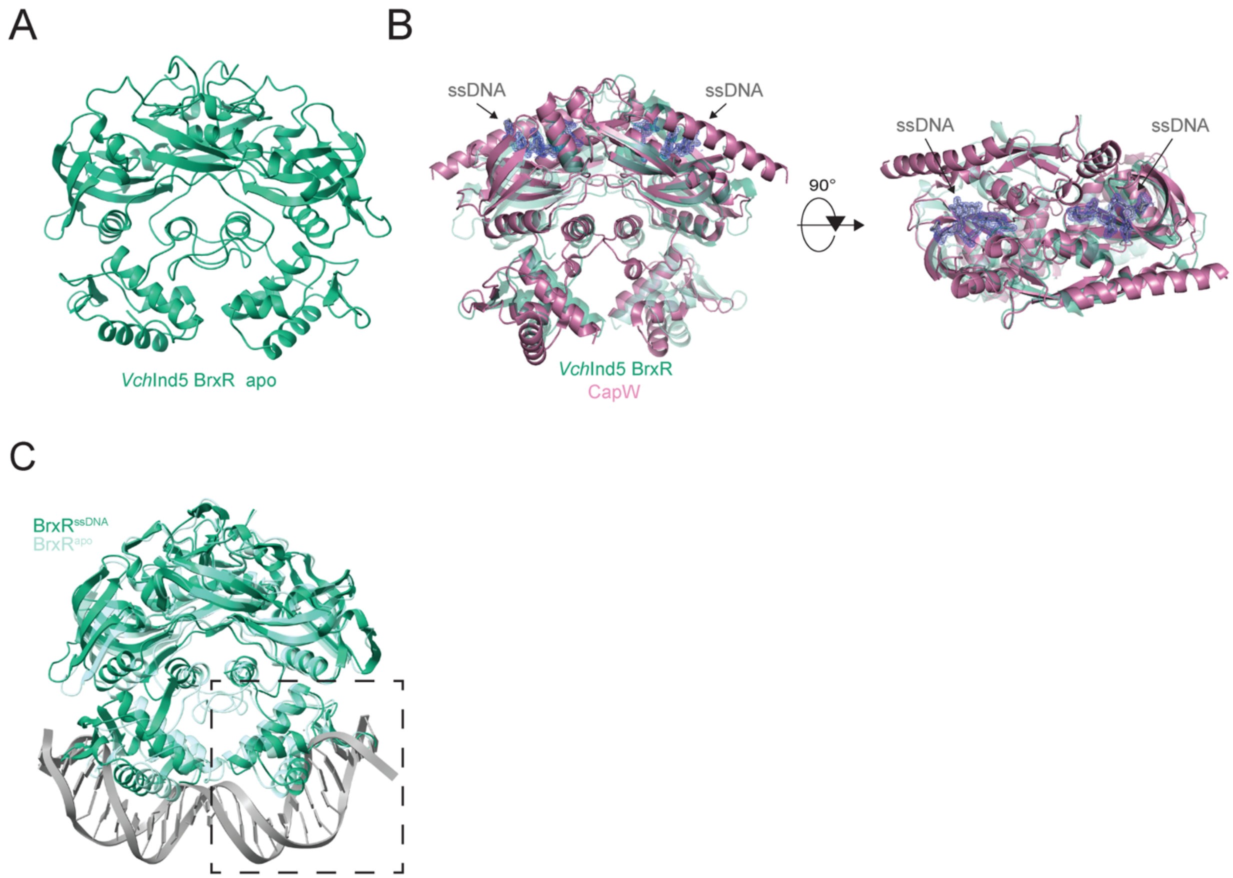


**Extended Data Figure 2. Single-stranded DNA binding leads to a conformational change in BrxR. A)** 3D crystal structure of BrxR encoded by the SXT ICE *Vch*Ind5 in the absence of single-stranded (ss) DNA. Crystallographic statistics are included in Supplementary Table 5. **B)** Comparison of the ssDNA-bound BrxR structure (light green) to the ssDNA-bound CapW (pink; PDB: 9C5G). **C)** Overlay of the apo and ssDNA-bound structure of BrxR overlaid with the *Acinetobacter* BrxR structure bound to double-stranded (ds) DNA (PDB: 7T8K). The dotted square indicates the zoomed-in image in Fig. 2E on the winged helix-turn-helix domain and the dsDNA to emphasize the clash that occurs when BrxR is bound to ssDNA. The Acinetobacter BrxR is not displayed.

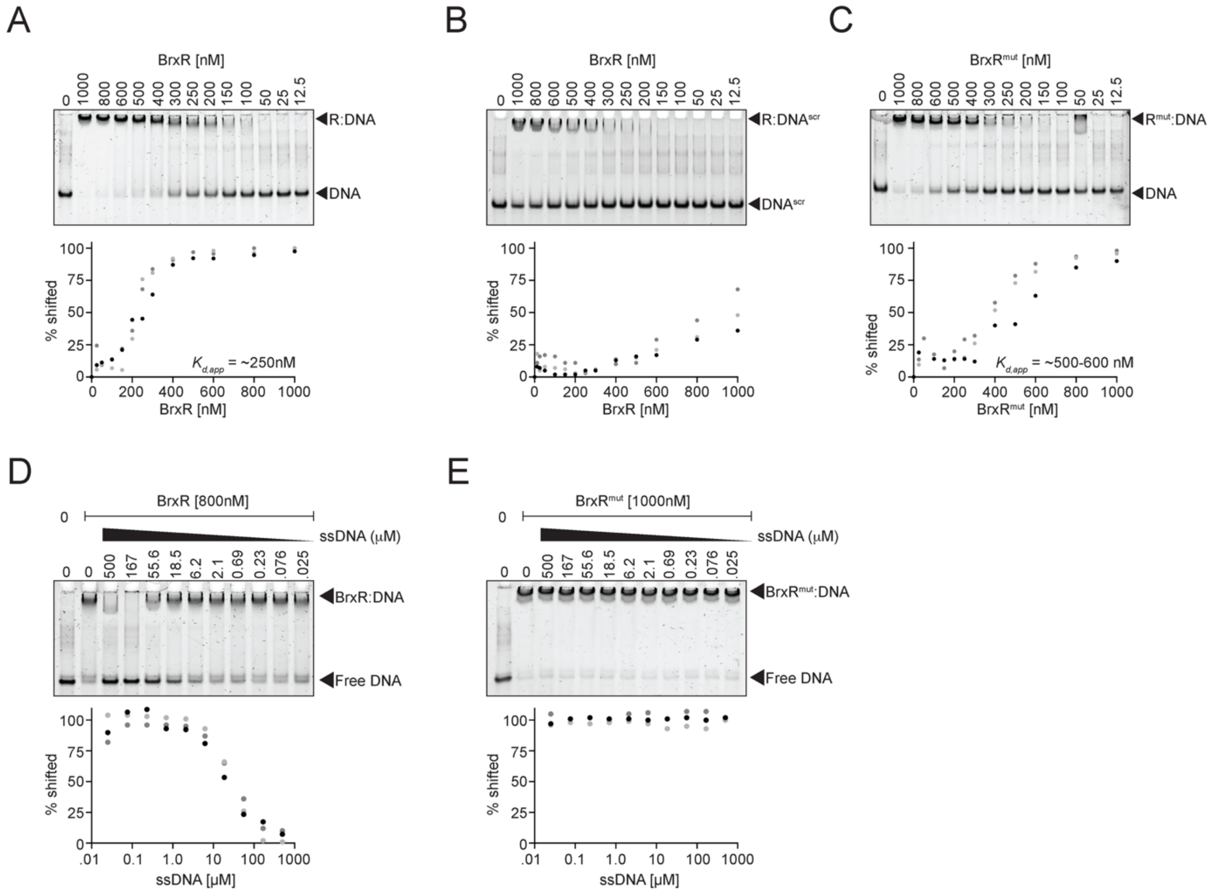


**Extended Data Figure 3. Mutation of BrxR's ssDNA binding pocket disrupts single-stranded DNA binding but slightly decreases double-stranded DNA binding affinity. A-C)** Electrophoretic mobility shift assays of either **A)** wild-type BrxR (R) with wild-type dsDNA target, **B)** wild-type BrxR with the dsDNA sequence scrambled (DNAscr), or C**)** the ssDNA-binding mutant ($R^{mut}$) bound to its wild-type dsDNA target. The concentrations of BrxR and $BrxR^{mut}$ proteins were titrated while the dsDNA concentration was held constant. The dsDNA target used harbors the BrxR inverted repeat sequence (see Methods for wild-type and scrambled sequences). Below are the associated binding curves for each replicate based on the densitometry analysis for the presented gel (black dots) and the replicates in Supplementary Figure 3 (dark and light gray dots). **D-E)** Electrophoretic mobility shift assays of either **D)** wild-type BrxR or **E)** the ssDNA-binding mutant ($BrxR^{mut}$) bound to its dsDNA target, being competed off with ssDNA. The concentration of BrxR and $BrxR^{mut}$ protein and its dsDNA target were constant while the concentration of ssDNA was titrated. The dsDNA target used harbors the BrxR inverted repeat sequence (see Methods for sequence). ssDNA used was a poly-T track (TTTTTT). Below are the calculated binding curves with ssDNA being titrated in the presence of either wild-type BrxR or the $BrxR^{mut}$. Binding curves were determined using gel images presented here and in Supplementary Figure 4.

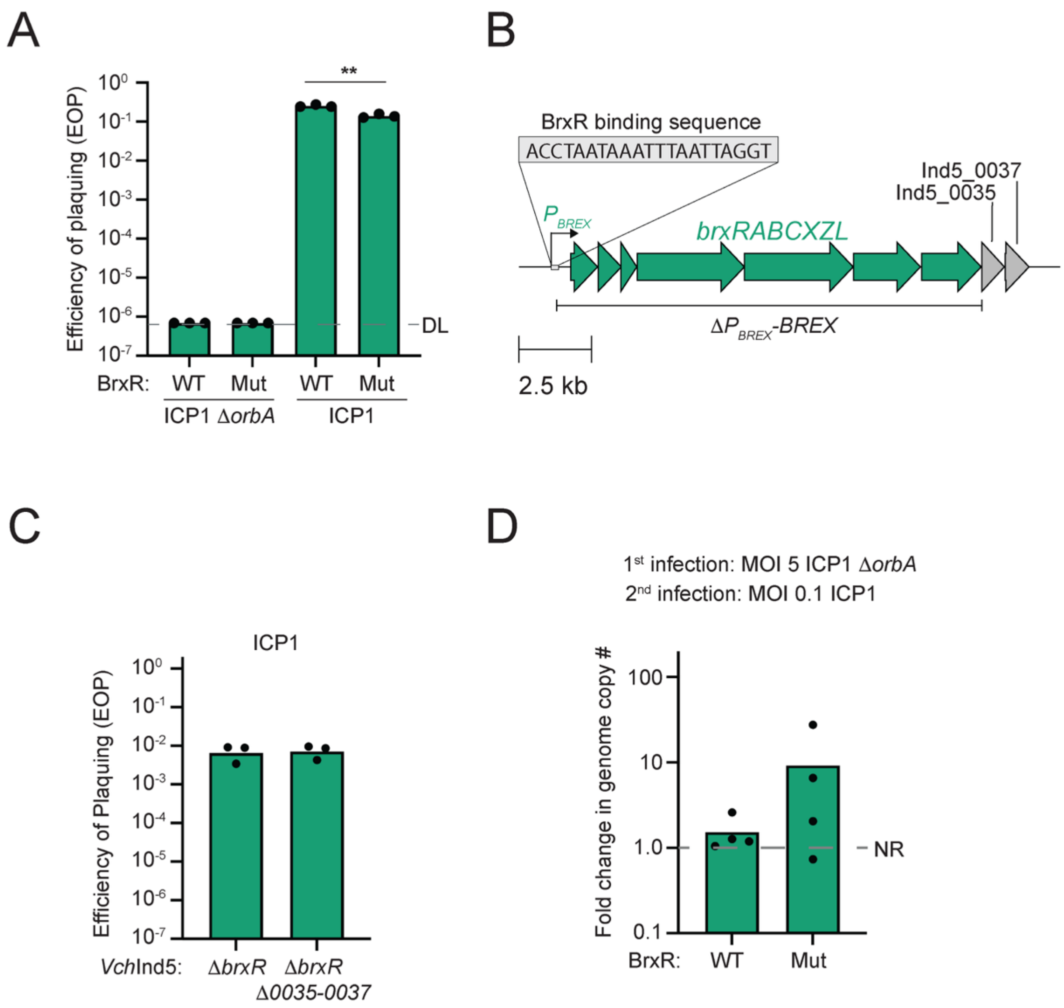


**Extended Data Figure 4. Increased BREX expression provides greater protection against phage-encoded BREX counterdefense. A)** Efficiency of plaquing (EOP) for both ICP1 Δ*orbA* and ICP1 in the wild-type BrxR background compared to the $BrxR^{mut}$ mutant background. DL indicates the detection limit. Statistical analysis was performed using a Student's t-test (** = $P$ < 0.005). Each dot represents an individual biological replicate. **B**) A gene map of the defense region in SXT *Vch*Ind5, depicting the BREX promoter ($P_{BREX}$), the BrxR-binding region and sequence, the BREX operon, the downstream genes (*Vch*Ind5_0035 and *Vch*Ind5_0037), and the boundaries for the $P_{BREX}$-BREX deletion. **C)** Efficiency of plaquing of ICP1 in the presence and absence of the two genes downstream of the BREX system (Panel B) in the *brxR*-deletion background. **D)** Quantification of ICP1 genome replication during a subsequent infection. Cells were first infected with ICP1 Δ*orbA,* and later infected with ICP1. The dashed line at one indicates there was no genome replication (NR). Each dot represents an individual biological replicate, and each bar represents the mean of the displayed biological replicates.

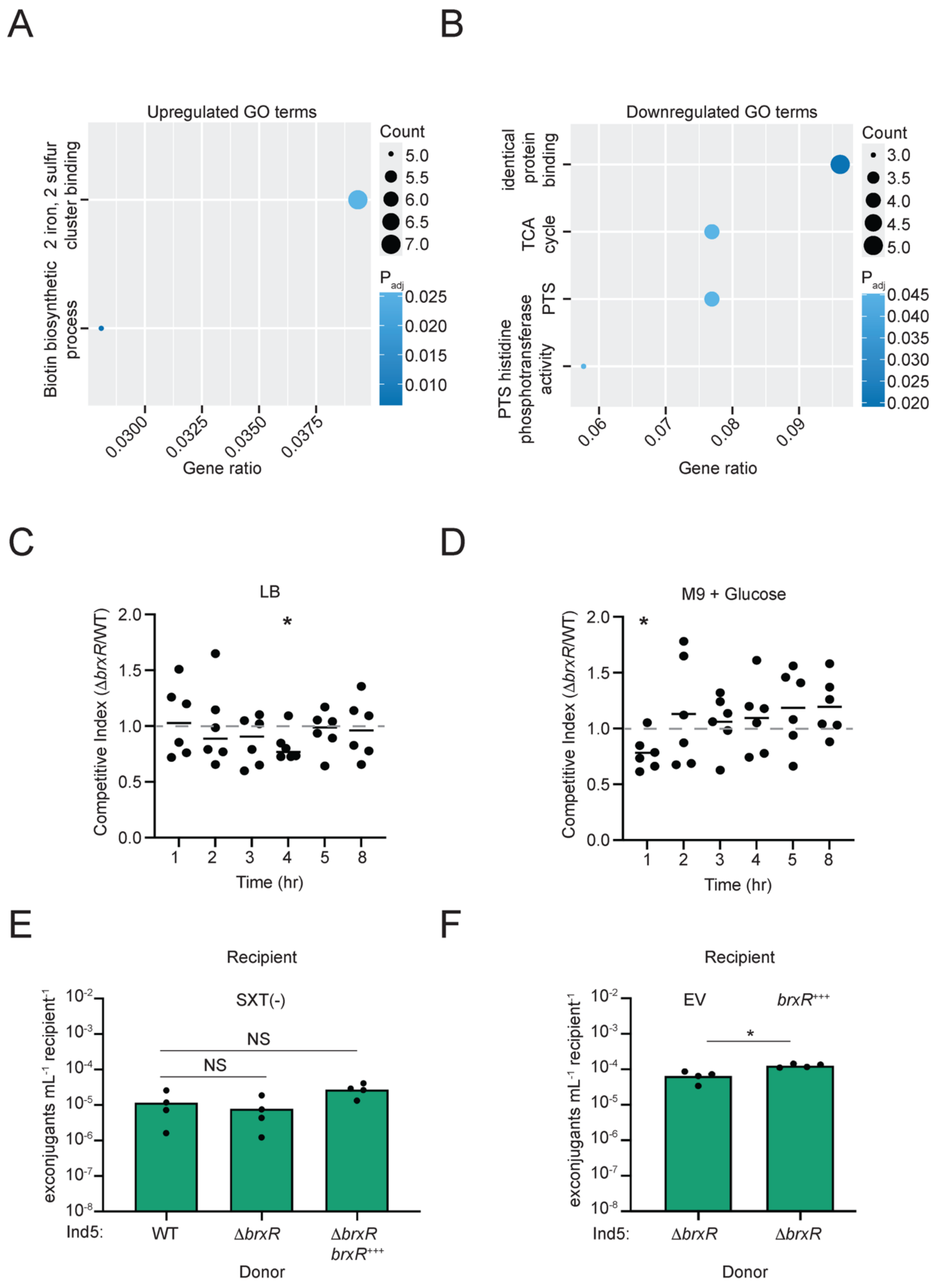


**Extended Data Figure 5. An increased abundance of BREX proteins in the absence of BrxR does not impact fitness or conjugation frequency. A-B)** Dot plots of gene ontology terms that were enriched from differentially expressed genes in the absence of BrxR versus the presence, that were either A) upregulated or B) downregulated. TCA cycle – tricarboxylic acid cycle. PTS – phosphoenolpyruvate-dependent sugar phosphotransferase system. PTS histidine phosphotransferase activity - protein-N(PI)-phosphohistidine-sugar phosphotransferase activity. For more details, see Supplementary Table 9. **C-D)** Competitive indices between Δ*brxR* and wild-type over time in **C)** LB or **D)** M9 + glucose. Dots represent individual biological replicates (n=6), and the line represents the mean. Statistical analysis was performed using a one-sample t-test and Wilcoxon test (* = $P$ < 0.05). **E-F)** Transfer efficiency of the SXT *Vch*Ind5 into recipient cells. **E)** Transfer efficiency into an SXT(-) recipient from three donor strains: wild-type (WT), Δ*brxR*, and the Δ*brxR* mutant expressing BrxR *in trans* (Δ*brxR* $brxR^{+++}$). Statistical analysis was performed using a one-way analysis of variance. NS, not significant. **F)** Transfer efficiency of Δ*brxR* *Vch*Ind5 donor into an SXT(-) recipient strain expressing either an empty vector (EV) or BrxR *in trans* ($brxR^{+++}$). Statistical analysis was performed using a Student's t-test (* = $P$ < 0.05). The bars represent the mean, and each dot represents a single biological replicate (n=4).

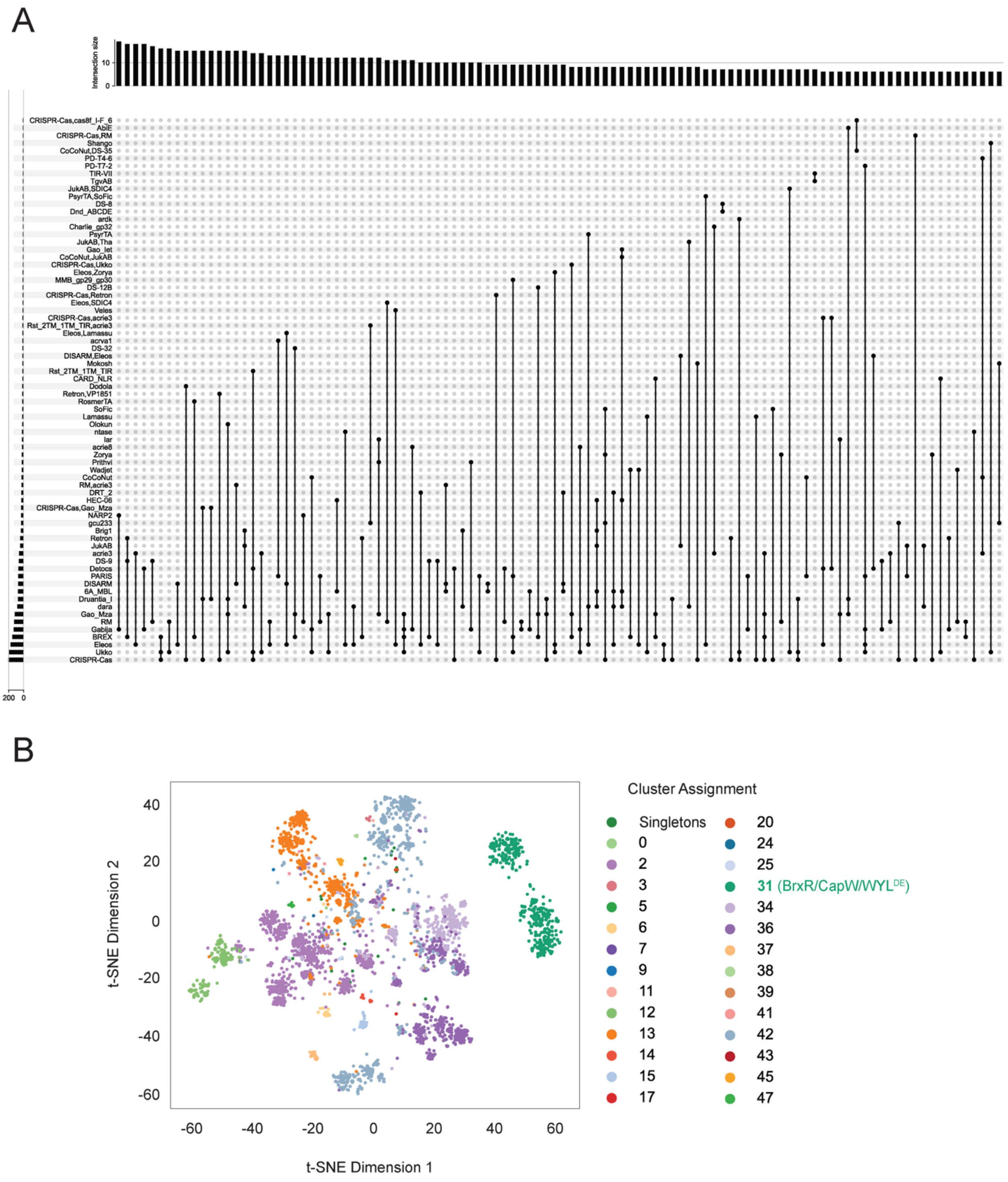


**Extended Figure 6. Clustering of defense-associated WYL proteins into distinct families. A)** UpSet plot showing the co-occurrence of defense systems that are associated with distinct daWYL proteins in the same genome, which are represented between 5 and 20 genomes. Defense systems that flank the same daWYL protein are listed as defense 1 and defense 2 (e.g., Eleos, Zorya). Intersection size indicates the number of genomes that represent a given combination. For the full list of defense combinations per genome, see Supplementary Table 14.

**B)** Scatter plot showing the relatedness of the defense-associated WYL proteins defined by Louvain community detection on pairwise Foldseek sequence alignments. BrxR/CapW/WYL$^{DE}$-like proteins are colored in green and represent Cluster 31. See Supplementary Table 15 for accession numbers associated with each cluster.

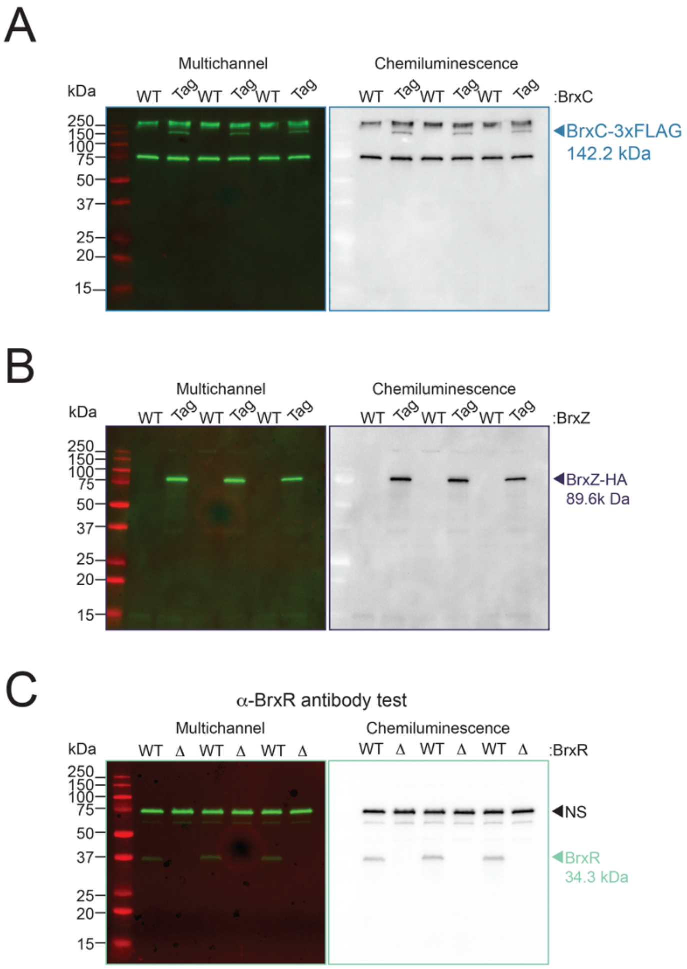


**Supplementary Figure 1. Testing untagged and tagged BREX proteins and the primary antibody against BrxR**. Uncropped replicate Western Blot images of **A)** Tagged BrxC, **B)** Tagged BrxZ, and **C)** the BrxR primary antibody. The band representing protein at its respective molecular weight is indicated by an arrow, along with its predicted molecular weight (kDa). WT – indicates the untagged BREX system. Tag – indicates that BrxC or BrxZ was tagged with 3xFLAG or HA, respectively. NS – nonspecific band that is used as a normalization control in the preceding experiments.

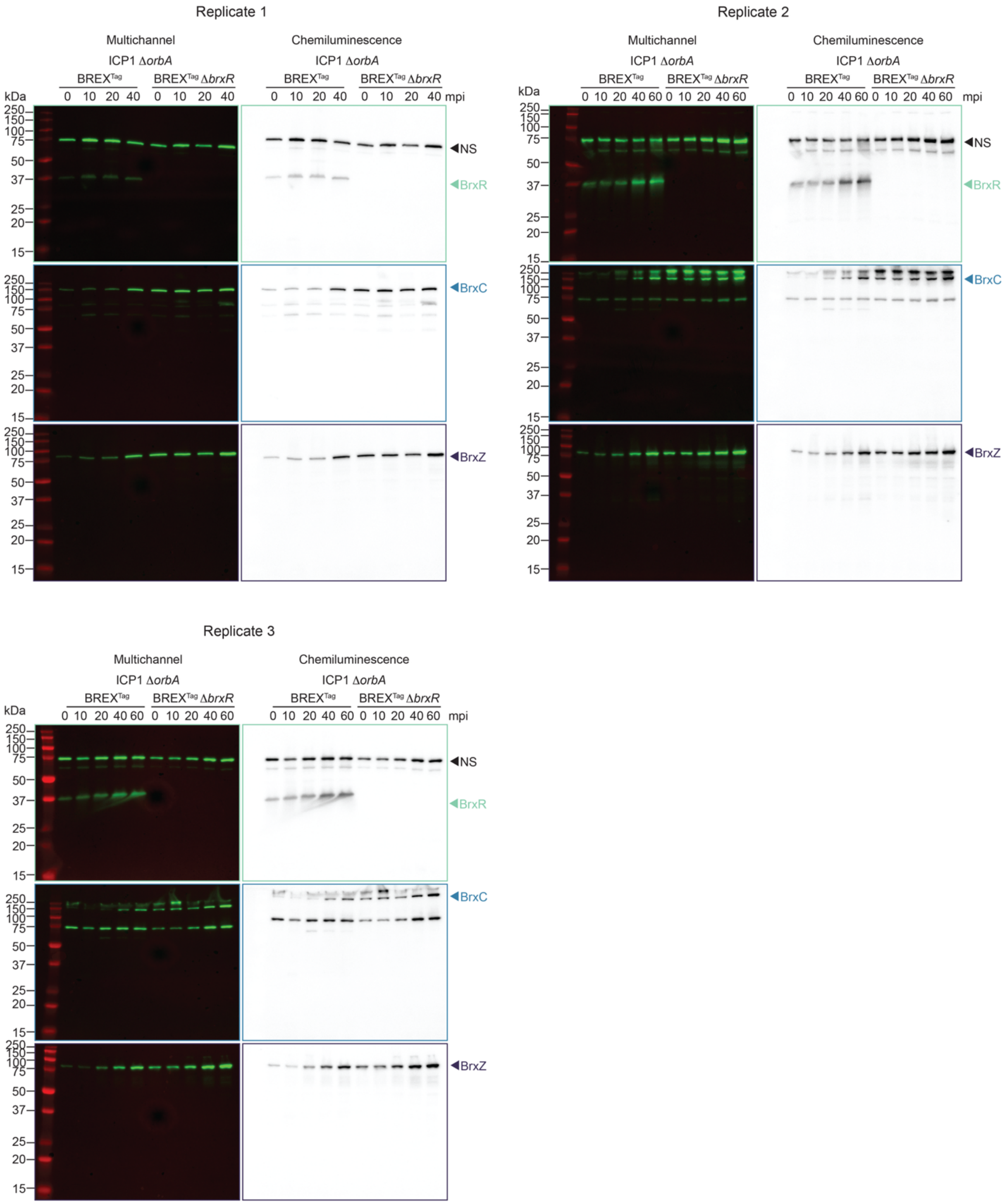


**Supplementary Figure 2. Induction of BREX during restricted phage infection is dependent on BrxR**. Uncropped replicate Western Blot images of phage infection with ICP1 Δ*orbA* at different time points (mpi, minutes post-infection) in the presence and absence of BrxR. Blots displayed in Figure 1C and Figure 2A are from replicate 1. The fold change calculation for Figure 1C and Extended Figure 2A was determined using the chemiluminescence gel. The multichannel gel is used to show the ladder and the corresponding molecular weights of each band. While a 60-minute time point is displayed, the data was not used for any calculations or discussed.

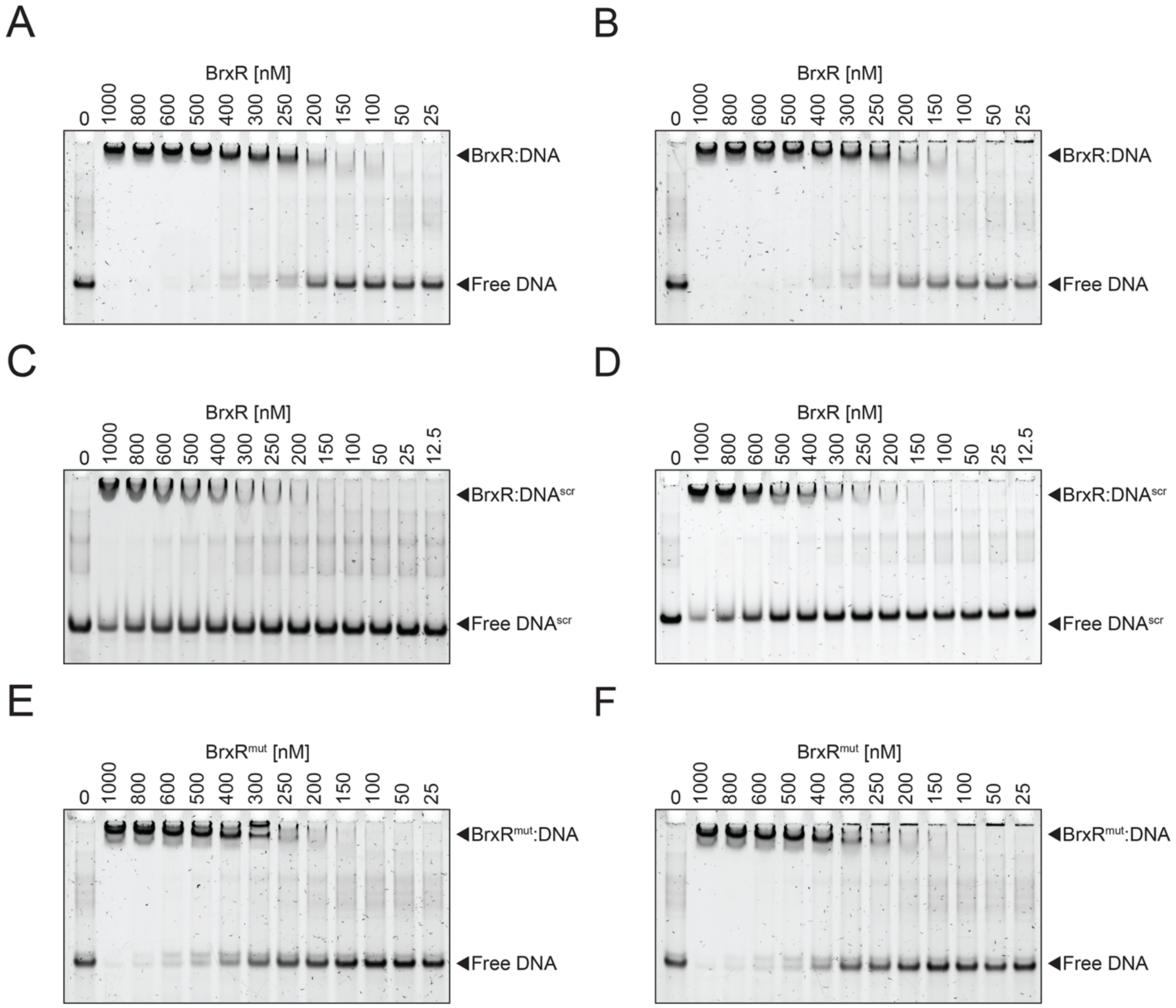


**Supplementary Figure 3. BrxR's ability to interact with ssDNA is not required for binding to dsDNA.** Electrophoretic mobility shift assays of either **A-B)** wild-type BrxR (R) with wild-type dsDNA target, **C-D)** wild-type BrxR with the dsDNA sequence scrambled (DNAscr), or **E-F)** the ssDNA-binding mutant ($R^{mut}$) bound to its wild-type dsDNA target. The concentrations of BrxR and $BrxR^{mut}$ proteins were titrated while the dsDNA concentration was held constant. The dsDNA target used harbors the BrxR inverted repeat sequence (see Methods for wild-type and scrambled sequences).

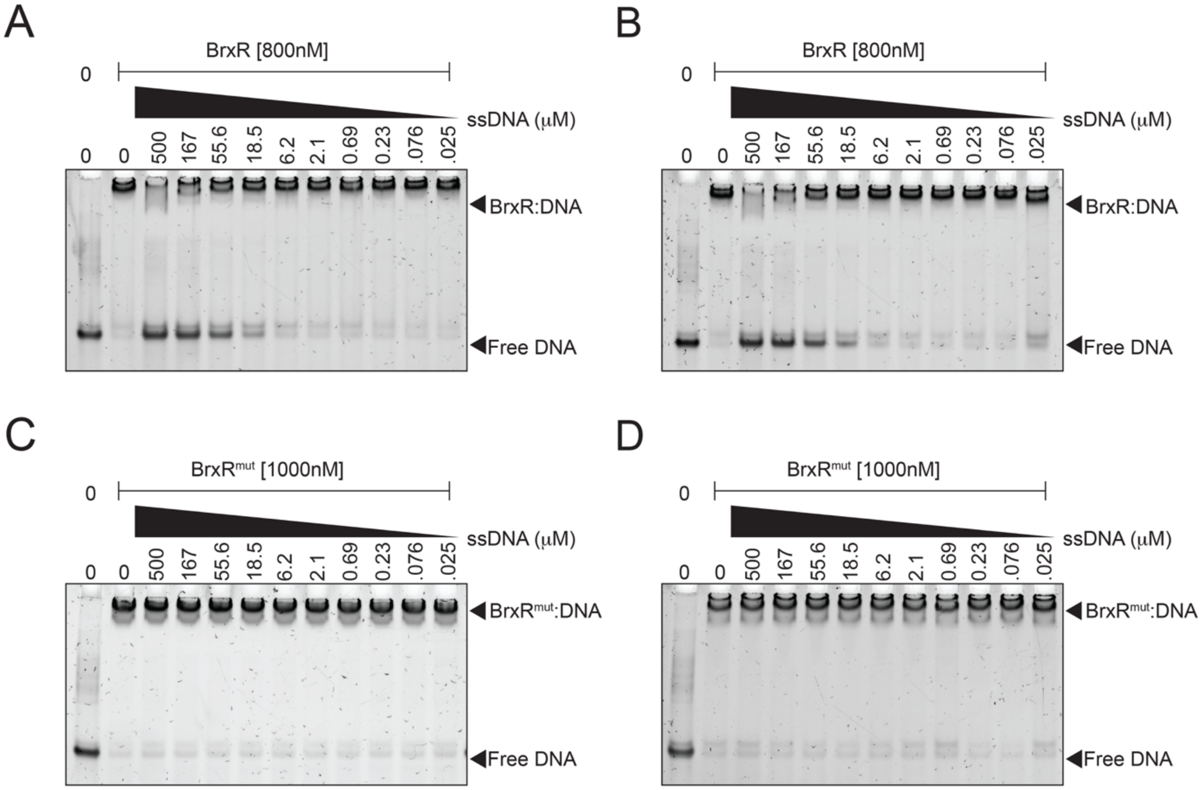


**Supplementary Figure 4. ssDNA binding is required for inhibition of BrxR's dsDNA-binding activity.** Electrophoretic mobility shift assays of either **A-B)** wild-type BrxR or **C-D)** the ssDNA-binding mutant (BrxR$^{mut}$) bound to its dsDNA target, being competed off with ssDNA. The concentration of BrxR and BrxR$^{mut}$ protein and its dsDNA target were constant while the concentration of ssDNA was titrated. The dsDNA target used harbors the BrxR inverted repeat sequence (see Methods for sequence). ssDNA used was a poly-T track (TTTTTT).

A

B

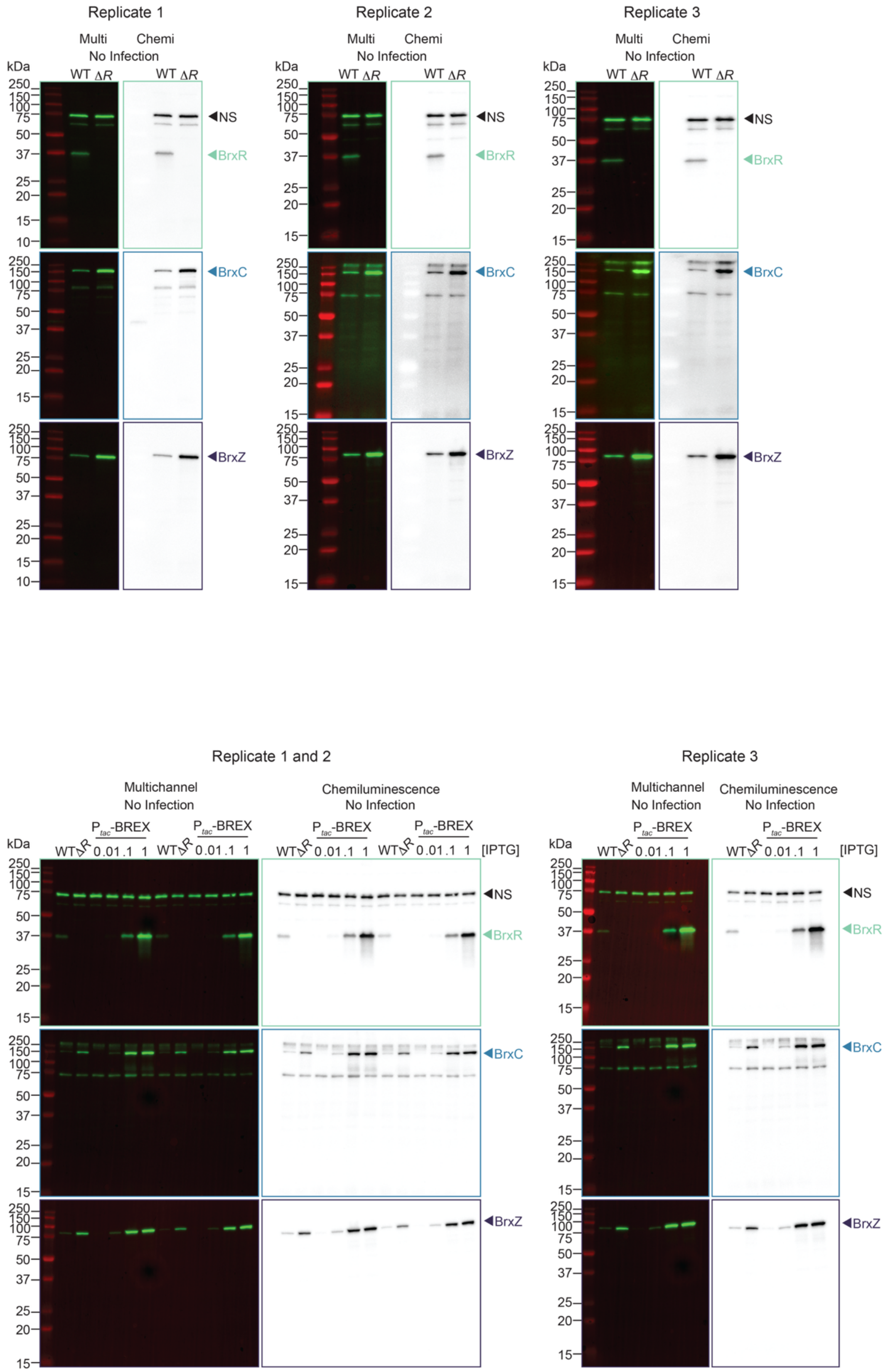
Replicate 1
Replicate 2
Replicate 3
Multi
Chemi
No Infection
kDa
WT ΔR
NS
BrxR
BrxC
BrxZ
250
150
100
75
50
37
25
20
15
10
Replicate 1 and 2
Multichannel
No Infection
Chemiluminescence
No Infection
P tac -BREX
WT ΔR 0 .01 .1 1
[IPTG]

**Supplementary Figure 5. BREX protein abundance can either be increased by the absence of BrxR or through an inducible promoter.** Uncropped replicate Western Blot images of **A)** the presence and absence of BrxR (ΔR), and **B)** induction of BREX ($P_{tac}$-BREX shortened from $P_{tac}$-riboE-BREX) using different amounts of inducer in comparison to the presence and absence of BrxR. The representative (chemi)luminescence images in Figure 3A are cropped from replicate 1. The representative images in Figure 3D are cropped from replicate 1 (left side lanes). The fold change calculation for Figure 3A and 3D was determined using the chemiluminescence gels displayed. The (multi)channel gel is used to show the ladder and the corresponding molecular weights of each band.
